# Strategies for enriching and characterizing proteins with inhibitory properties on the yeast surface

**DOI:** 10.1101/2022.04.20.488975

**Authors:** Arlinda Rezhdo, Catherine T. Lessard, Mariha Islam, James A. Van Deventer

## Abstract

Display technologies are powerful tools for discovering antibodies and other binding proteins against a broad range of biological targets. However, it remains challenging to adapt display technologies for the discovery of proteins that inhibit the enzymatic activities of such targets because the phenotypic readout during display screens is binding. The goal of this work is to investigate approaches for discovering inhibitory antibodies in yeast display format using a well-defined series of constructs and the target matrix metalloproteinase-9 (MMP-9). Three previously reported antibodies (DX-2802, M0076 and FAPB2.3.6) were used to create model libraries that are representative of protein libraries consisting of inhibitory binders, non-inhibitory binders, and non-binding constructs. Conditions that preferentially enrich for inhibitory clones were identified for both magnetic bead-based enrichments and fluorescence-activated cell sorting (FACS). Finally, we used direct titration of yeast to estimate inhibitor IC_50_ values with yeast-displayed and soluble constructs and found that the IC_50_ obtained for DX-2802 in yeast display format (20.01 ± 9.01 nM) falls within the confidence interval of IC_50_ the soluble scFv-Fc form of DX-2802 (17.56 ± 6.16 nM). Thus, it is possible to obtain IC_50_ values on the yeast surface, which greatly streamlines initial characterizations of inhibitory properties. Overall, we used these well-defined constructs to identify strategies for the discovery and characterization of inhibitory clones directly in surface display format.

## Introduction

Display technologies—including yeast display, phage display, mRNA display, and others—are critical tools for isolating binding proteins and peptides with desired properties^1–5^. These platforms support discovery of affinity ligands against diverse biological targets, engineering of existing ligands for improved binding affinity and stability^6, 7^, profiling proteinprotein interactions^8^, advancing bioorthogonal reactions^9^ and more^10^. Display-based selection and screening platforms exhibiting a wide array of properties are now available, ranging from entirely cell-free platforms (e.g., mRNA display), to platforms based on phage or viruses (e.g. phage display and virion display), to cell surface display platforms derived from the cells of organisms ranging from Gram-positive and Gram-negative bacteria to yeast, insects^11^ and humans^2, 12^. Arguably, the rise of this versatile set of technologies has made the discovery of high affinity binders against many biological targets routine^3, 4, 13, 14^. However, simply binding to a target is not always sufficient to modulate key target functions. This is particularly true in the case of enzymes, many of which remain challenging to inhibit specifically. Although display platforms are not necessarily well-suited for the discovery of enzyme inhibitors, careful adaptations of display techniques provide flexibility for identifying and characterizing protein-based enzyme inhibitors.

The key challenge in utilizing display technologies for identifying enzyme inhibition is that display technologies utilize binding events, not inhibition, as the primary phenotype during enrichments and characterizations. Previous adaptations of display technologies have made it possible to evaluate numerous protein properties in display format, but strategies linked to the inhibitory properties of displayed proteins remain limited in scope. Substantial efforts to tailor the specificities of naturally occurring, protein-based inhibitors, including tissue inhibitors of metalloproteinases (TIMPs), have utilized competitive binding assays in display format to distinguish between on- and off-target binding^15, 16^. This competition strategy has also been utilized to evaluate whether inhibitory antibodies compete for binding with TIMPs on the yeast surface^17–20^. Selections of inhibitory antibody fragments from naïve antibody libraries have been described utilizing phage display^21^ and selections in the periplasm of *E. coli*^17^ using competitions with small-molecule inhibitors and a selection marker for protease cleavage, respectively. These powerful approaches demonstrate the feasibility of identifying inhibitory proteins during discovery campaigns and characterizations in multiple formats. However, these efforts also highlight substantial opportunities to further extend and enhance these approaches. In particular, the quantitative enrichments and characterizations supported by yeast display merit further exploration to address the unique challenges posed by the discovery of protein-based inhibitors.

Here, we investigate strategies for adapting yeast display to the discovery and characterization of protein constructs with inhibitory properties (Fig. 1). We focused on devising enrichment and characterization strategies by utilizing previously reported antibodies that target matrix metalloproteinase-9 (MMP-9) in multiple ways: an inhibitory antibody to MMP-9 (DX-2802)^22, 23^, an antibody that binds to MMP-9 with no detectable inhibition (non-inhibitory clone M0076-D03)^22, 23^, and a control antibody that does not bind to MMP-9 (FAPB2.3.6)^24^. Flow cytometric analyses with yeast-displayed forms of these antibodies revealed that each of these constructs exhibits distinct display and MMP-9 binding properties following incubation with active MMP-9. With the unique signatures associated with the three clones, we were able to utilize flow cytometry analysis to investigate a number of enrichment schemes using magnetic beads and fluorescence-activated cell sorting both qualitatively and quantitatively. To evaluate enrichment strategies, we combined the constructs in defined ratios and investigated multiple schemes to identify schemes for preferentially isolating inhibitors. For bead-based enrichments, we compared standard enrichments using MMP-9 immobilized on beads to a strategy in which MMP-9 was incubated with the yeast population prior to the introduction of magnetic beads. This solution-based preincubation step resulted substantially increased enrichment ratios for inhibitory binders compared to the enrichment ratios obtained with standard bead-based sorting conditions. Similarly, fluorescence-activated cell sorting results showed that sorting using a “high gate” leads to efficient enrichment inhibitory binders, even starting from a population in which inhibitory clones were far outnumbered by non-inhibitory clones. Finally, we evaluated half maximal inhibitory concentrations (IC_50_s) directly on the yeast surface and compared these values to solution-based determinations of IC_50_ values. Measurements made by titrating increasing numbers of yeast-displaying antibodies into wells containing a fixed concentration of MMP-9 led to IC_50_ values that were indistinguishable from IC_50_s obtained with a soluble construct of the inhibitory antibody, indicating that this parameter can be determined without protein purification. The studies conducted with the model system described in this work establish promising approaches for pursuing more efficient discovery and characterization of protein-based inhibitors directly in yeast display format.

**Fig. 1.**
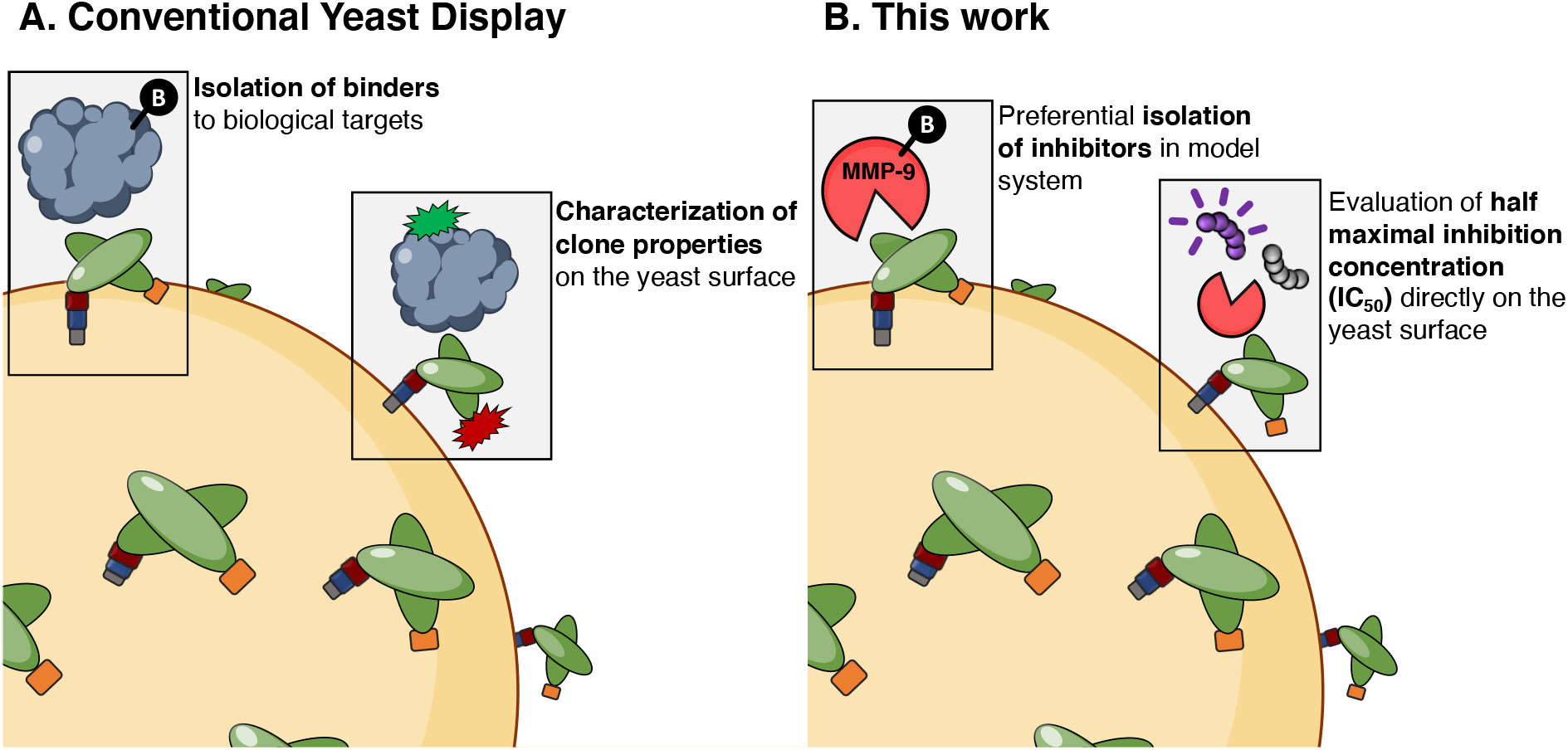
Schematic of yeast display and specific investigations conducted in this work. (**A**) Yeast display is most commonly used for isolation of binders to biological targets and facilitates general characterization of clone properties including affinity and stability. (**B**) This work evaluates strategies for the preferential isolation of inhibitors and the determination of half maximal inhibition concentration (IC_50_) in yeast display format. These investigations use a series of individual clones and the model target matrix metalloproteinase-9 (MMP-9).

## Results

### Establishment of model system

To establish a model system for evaluating the feasibility of isolating protein inhibitors displayed on the yeast surface, we prepared yeast displayed versions of three previously described constructs in scFv format. The antibody constructs used here are DX-2802, a known inhibitor of MMP-9^22, 23^, M0076-D03 (referred to here as M0076), a non-inhibitory MMP-9 binder^22, 23^, and FAPB2.3.6, which recognizes the unrelated target fibroblast activation protein^24^ but does not bind to or inhibit MMP-9. All sequences are available in Supplementary Table S1. These constructs comprise three major types of clones expected to be present in diverse antibody libraries: inhibitory binders, non-inhibitory binders, and clones that do not recognize the target antigen.

To evaluate the binding and display properties of each clone in yeast display format, we conducted flow cytometry binding assays where each of the constructs displayed on the yeast surface was incubated with the pro form of human MMP-9 (pro-hMMP-9), trypsin-activated hMMP-9 (act-hMMP-9), or deactivated hMMP-9 prepared by incubating activated MMP-9 with the chelator EDTA (deact-hMMP-9; see *Materials and Methods* for details on preparation of MMP-9 samples). Following incubation, the yeast samples were labeled for detection of MMP-9 binding and full-length antibody display. Two-dimensional dot plots (Fig. 2A, B) and quantitative analyses of fluorescence levels (Fig. 2C, D) reveal large differences in behavior between the three constructs. As expected, FAPB2.3.6, the non-binding control, shows no detectable levels of binding to pro-, act-, or deact-hMMP-9, and unchanged display levels following incubation with any form of MMP-9. DX-2802, the inhibitory binder, exhibits high levels of hMMP-9 detection only after incubation with act-hMMP-9, and exhibits no detectable binding to pro- or deact-hMMP-9. M0076, the non-inhibitory binder, exhibits binding to pro-hMMP-9 but essentially no binding to the other two forms of hMMP-9 investigated here. Interestingly, incubating act-hMMP-9 with yeast displaying M0076 resulted in decreased levels of cMyc detection in comparison to cMyc levels following incubation with no hMMP-9 (control) or with pro- or deact-hMMP-9 (Fig. 2D). Given the high enzymatic activity of act-hMMP-9, and due to the presumed proximity of its active site to the displayed form of M0076 following binding, we hypothesize that the reduction in cMyc signal is due to MMP-9-mediated cleavage of the M0076 display construct at a location prior to the cMyc epitope tag. This cleavage appears to be mediated by the binding of MMP-9 to M0076 prior to the cleavage event. This hypothesis is consistent with our observation that there is no detectable reduction in cMyc signal following incubation of any form of hMMP-9 with either DX-2802 or FAPB2.3.6 (Supplementary Fig. S1). Taken together, the three constructs utilized here exhibit MMP-9 binding levels and construct display levels that are readily distinguishable from one another. Thus, they form a well-behaved model system for investigating selections, screens, and characterization strategies for the preferential identification and enrichment of inhibitory clones.

**Fig. 2.**
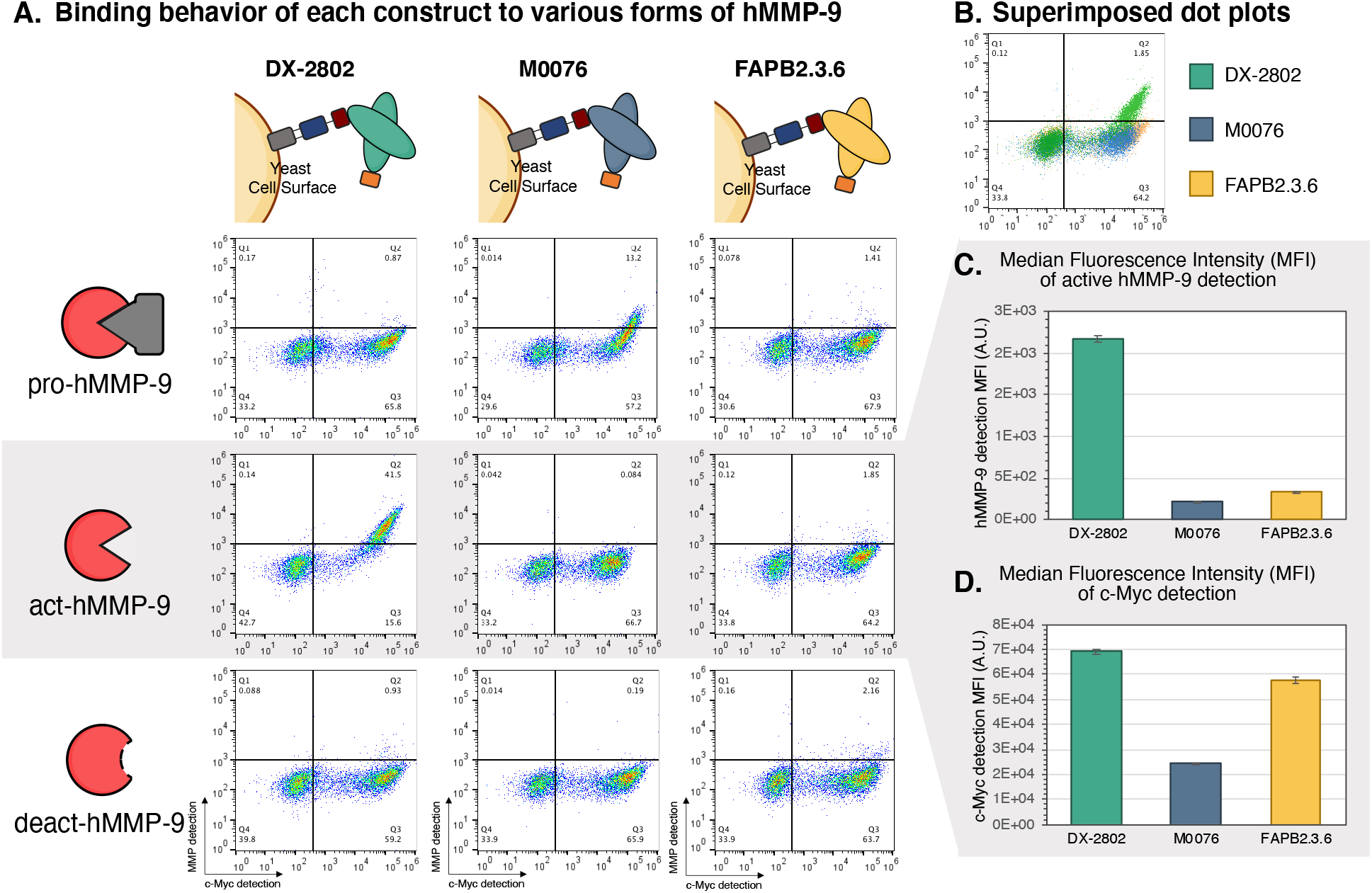
Analysis of the binding behaviors of DX-2802, M0076, and FAPB2.3.6 on the yeast surface with various forms of hMMP-9. (**A**) Flow cytometry dot plots of yeast surface displayed DX-2802, M0076, and FAPB2.3.6 when incubated with 200 nM of pro-hMMP-9 (inactive), act-hMMP-9 (activated with trypsin), and deact-hMMP-9 (deactivated with EDTA) shows observable differences in MMP-9 binding levels and cMyc detection levels. (**B**) Superimposed flow cytometry dot plots of the three displayed clones after treatment with 200 nM act-hMMP-9. (**C**) Median fluorescence intensity (MFI) of act-hMMP-9 binding levels for all three constructs. (**D**) Median fluorescence intensity (MFI) of cMyc detection for all three constructs following incubation with 200 nM act-hMMP-9.

### Magnetic bead-based enrichments

With yeast display-based libraries, immobilization of antigens on magnetic beads readily supports enrichments with 10 billion or more cells on the benchtop. Here, we sought to investigate whether we could identify enrichment conditions that would selectively enrich yeast cells displaying inhibitors in preference to cells displaying binders. To evaluate different enrichment schemes, we prepared four “model libraries” with varying ratios of the three described constructs. Model libraries containing mixtures of DX-2802, M0076 and FAPB2.3.6 with the ratios (i) 1:1:100, (ii) 1:100:100, (iii) 1:1:1000, and (iv) 1:1000:1000 were prepared as described in *Materials and Methods.* The libraries were enriched using two main conditions: (1) *antigen-on-beads condition* – cells were incubated with antigen-coated beads; and (2) *antigen-in-solution condition* – cells were incubated with antigen in solution, washed, and then incubated with bare streptavidin-coated beads (see more details in *Materials and Methods*). In all cases, the antigen was active, biotinylated hMMP-9, and the samples were all handled identically during magnetic selections and subsequent washing steps.

To evaluate enrichment efficiency, we conducted flow cytometry characterizations for each library before and after magnetic selections (Fig. 3). Single clones of DX-2802, M0076 and FAPB2.3.6 were first analyzed and gated. These same gates were applied to the flow cytometry plots of each of the mixtures before and after magnetic selections and were used to quantitatively determine fold enrichment (Supplementary Fig. S2). Pre-enrichment evaluation of hMMP-9 binding and cMyc detection levels confirmed that the estimated composition of constructs in each model library matched the intended compositions of interest (Fig. 3A). Flow cytometric analysis following enrichment shows clear changes in the compositions of the populations using either the *antigen-on-beads* condition or *antigen-in-solution* condition (Fig. 3, Supplementary Fig. S2). Under the *antigen-in-solution* condition, the increased number of events exhibiting both act-hMMP-9 binding and cMyc positivity strongly indicates the preferential enrichment of yeast displaying DX-2802 (Fig. 3A). Under the *antigen-on-beads* condition, the flow cytometry dot plots indicate only low levels of act-hMMP-9 binding and a decrease in cMyc display levels, suggesting that cells displaying the M0076 non-inhibitory clone are being enriched under these conditions (Fig. 3A). Thus, qualitative analysis indicated readily discernable changes in the compositions of model libraries after a single round of enrichment, with construct preference being dependent on the type of enrichment used.

**Fig. 3.**
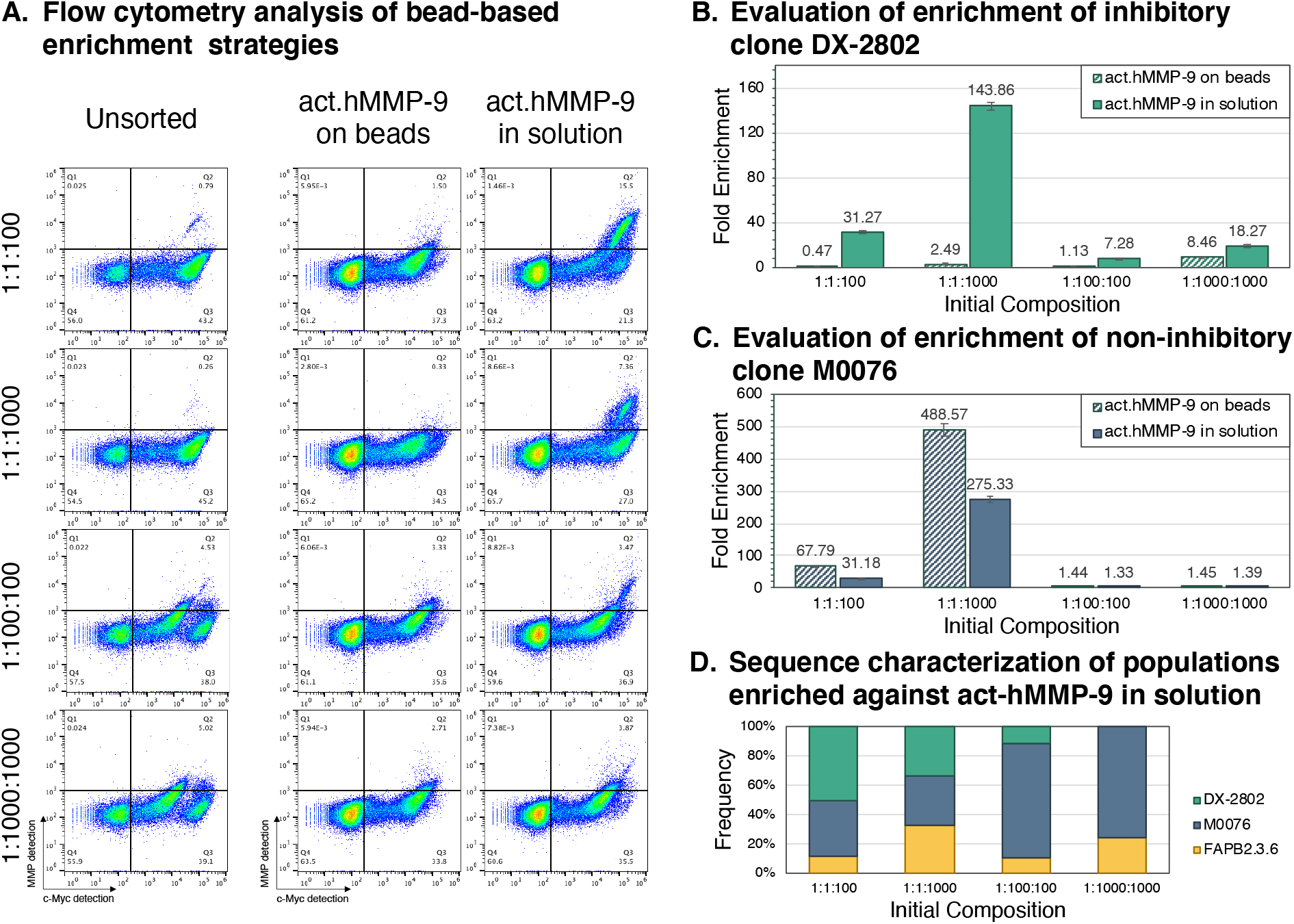
Evaluation of bead-based enrichment strategies. (**A**) Flow-cytometry plots of sorting results for four different compositions enriched in two different conditions. Analysis of bead-based enrichments for (**B**) inhibitors and (**C**) binders for both conditions of MMP activated on beads, and MMP activated and incubation in solution. All data represented here was performed in technical triplicates. (**D**) Sequence characterization of ten randomly picked colonies of each population sorted with active hMMP-9 in solution.

To further evaluate the outcomes of these experiments, we performed quantitative flow cytometric analyses of enrichment outputs based on the distinct binding properties of the three clones. To do this, we defined separate gates drawn in the regions where the majority of DX-2802 and M0076 clones appear in two-dimensional dot plots (Supplementary Fig. S2). The events appearing in each gate were compared to the total number of cMyc-positive events to obtain the fraction of each clone appearing in a given population. “Fold enrichment” was calculated for all enriched samples by taking the fraction of events appearing in gates of interest in the enriched populations and dividing these values by the fraction of events present in the gate in the initial population (Fig. 3B; see Supplementary Fig. S2 for gating) using the two equations below:

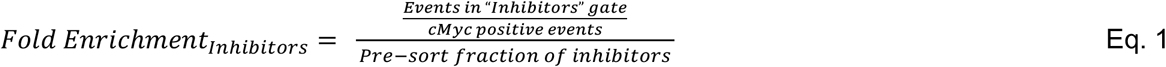

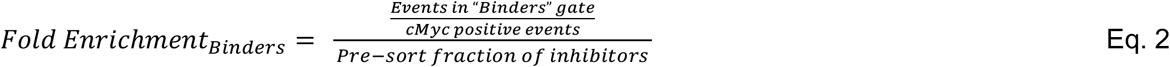

Notably, under the *antigen-in solution* condition, DX-2802 was enriched by 31.3-fold and 144-fold in the 1:1:100 and 1:1:1000 compositions respectively, compared to 0.47-fold and 2.49-fold when the *antigen-on-beads* strategy was utilized. In the population compositions where inhibitory DX-2802 clones are far outnumbered by M0076 clones (1:100:100 and 1:1000:1000), DX-2802 was enriched by 7.28-fold and 18.3-fold respectively, compared to 1.13-fold and 8.46-fold when enriched utilizing the *antigen-on-beads* strategy (Fig. 3B). In all cases the enrichment of the inhibitor is significantly higher under the *antigen-in-solution* condition compared to the *antigen-on-beads* condition. Enrichment of M0076 is more prevalent for the *antigen-on-beads* condition, with values being as high as 67.9-fold and 489-fold in the 1:1:100 and 1:1:1000 compositions, respectively, compared to 31.2-fold and 275-fold when enriched with the *antigen-in solution* condition. For three out of four ratios investigated here, the enrichment of inhibitory clones under the *antigen-in-solution* condition is higher than under the *antigen-on-beads* condition, combined with a decrease in the enrichment for non-inhibitory binders, signifying a preferential enrichment of inhibitors under the *antigen-in-solution* condition. To confirm the trends observed via flow cytometry, we isolated individual clones from one replicate of each enriched population and determined the sequences of 9–10 clones per condition (Fig. 3D; see *Materials and Methods* for details). This sequencing revealed that clones were present in the enriched populations at frequencies consistent with the frequencies determined via flow cytometric analysis. Overall, these results show that steps or conditions of bead-based enrichment protocols can be tested and adjusted to support preferential enrichment of inhibitors over binders, even when the binding clones of a population are initially present in vast excess.

### Flow-cytometry based enrichments

In most yeast display campaigns, fluorescence activated cell sorting (FACS) is performed in later rounds to isolate clones exhibiting the most promising properties of interest^25–27^. Here, we sought to determine whether we could preferentially isolate clones with inhibitory properties while discriminating against binders without such properties. Our bead-based enrichments (Fig. 3) indicated that isolation of inhibitors appears to be more challenging using the ratios 1:100:100 and 1:1000:1000, where cells displaying the binding protein M0076 vastly outnumber cells displaying DX-2802. Therefore, to determine the efficiency of FACS-based isolations of inhibitors, we prepared samples of 10^7^ cells containing the 1:100:100 and 1:1000:1000 ratios (see *Materials and Methods* for details). Following incubation with active hMMP-9 for 15 minutes, library populations were sorted via FACS based on their relative levels of cMyc and hMMP-9 detection (Fig. 4). Two gates were drawn to isolate distinct populations of cells: The “high gate” was drawn to isolate cells exhibiting high levels of both cMyc and hMMP-9 binding, while the “low gate” was drawn to isolate clones exhibiting moderate levels of cMyc and background levels of hMMP-9 binding (Supplementary Fig. S3).

**Fig. 4.**
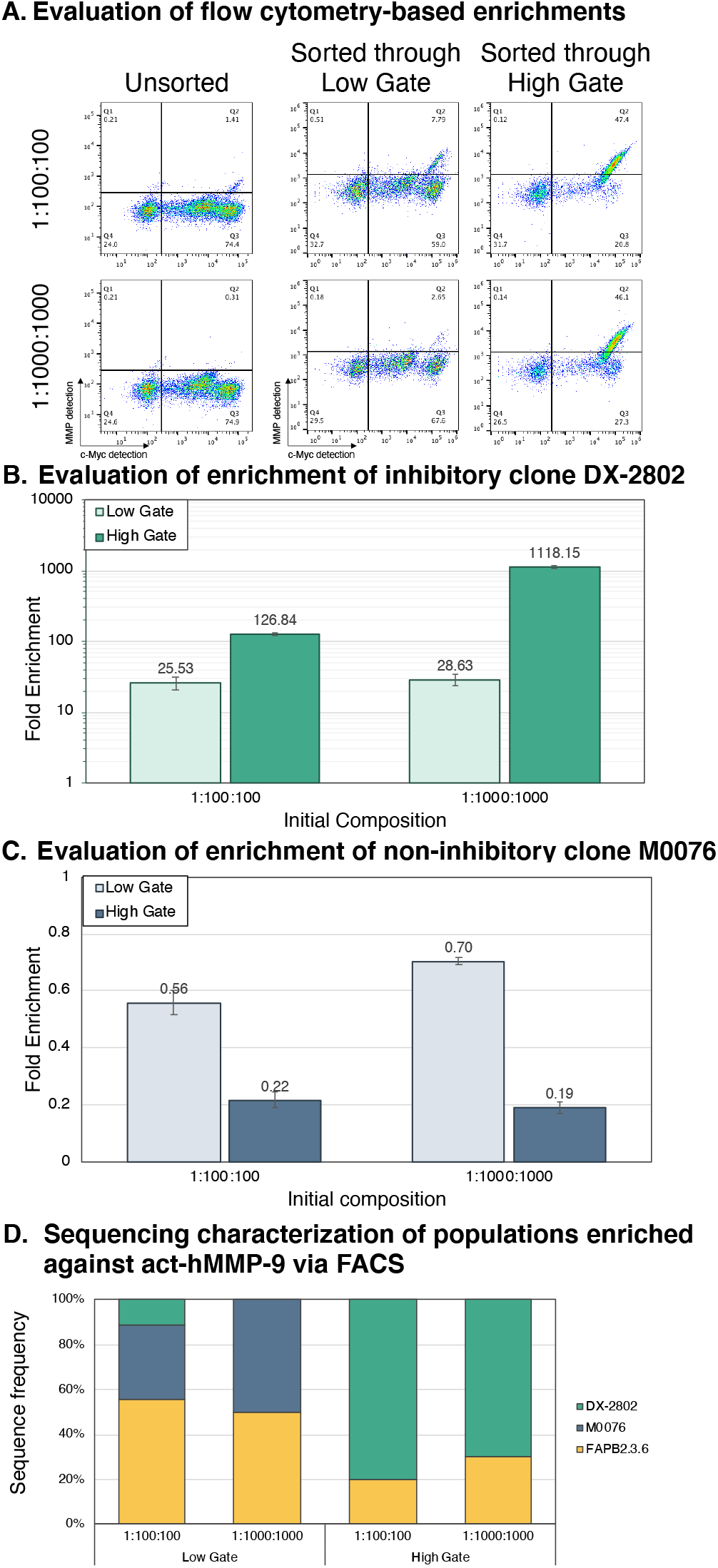
Evaluation of fluorescence-activated cell sorting to preferentially enrich inhibitors. (**A**) Flow cytometry-based enrichments using a low and high gate for two different populations. Analysis of flow cytometry-based enrichments of (**B**) inhibitors and (**C**) binders for both low and high gates. All experiments were performed in technical triplicate. (**D**) Sequence characterization of ten randomly picked individual colonies from one replicate of each sorted population.

Following recovery of sorted populations, we evaluated the act-hMMP-9 binding levels and cMyc display levels of all populations. Two-dimensional flow cytometry dot plots show clear differences between populations isolated via the high and low gates for both ratios (Fig. 4B, see Supplementary Fig. S4 for triplicate data). The substantial proportion of cells exhibiting both hMMP-9 binding and high cMyc levels strongly suggests the enrichment of inhibitory binders. To quantitatively evaluate enrichment efficiencies, we again defined gates intended to capture inhibitory binders and non-inhibitory binders and determined fold enrichment as for the bead-based enrichment experiments (Supplementary Fig. S4). Populations isolated via the high gate exhibit higher numbers of events that correspond to inhibitors compared to the populations isolated via the low gate (Fig. 4C). In the 1:100:100 composition, sorting via the high gate results in almost 5 times higher enrichment for inhibitors compared to the low gate, while in the 1:1000:1000 composition, sorting via the high gate results in almost 40 times higher enrichment for inhibitors compared to the low gate. To further confirm these results, we again isolated DNA from one set of replicates of the enriched populations and sent in individual clones for sequencing (see *Materials and Methods* for details). Sequences of 10 randomly chosen clones revealed frequencies consistent with the frequencies determined by flow cytometry-based analysis (Fig. 4D). Taken as a whole, these results indicate that inhibitory clones can be preferentially enriched over binding clones in FACS even when binding clones are present in vast excess.

### Evaluating IC_50_ of constructs on the yeast surface

Evaluating the properties of individual clones is a critical part of identifying lead clones with desirable properties. However, the extent to which inhibitory properties can be measured on the yeast surface remains unclear. Here, we conducted assays comparing IC_50_ value determination using yeast displayed constructs and soluble constructs to investigate whether yeast display supports the same level of quantitation of inhibition as solution-based measurements. Fig. 5A depicts schematics of the yeast display assay and time-dependent fluorescence data collected for titrations with increasing numbers of yeast. We define effective scFv concentration as the concentration of scFvs in the sample as if they were free-floating in solution (i.e. not bound to cells). Assuming 5×10^4^ constructs are displayed on a fully induced yeast cell ^28^, we titrated numbers of cells corresponding to the effective concentrations listed in the figure into individual wells containing 2 nM active hMMP-9 (Fig. 5A). Addition of FS-6, a fluorogenic substrate of MMP-9, to each well and monitoring fluorescence over time revealed that increasing numbers of cells displaying DX-2802 led to decreased rates of fluorescence evolution. In contrast, addition of increasing numbers of cells displaying either M0076 or FAPB2.3.6 did not affect fluorescence evolution.

**Fig. 5.**
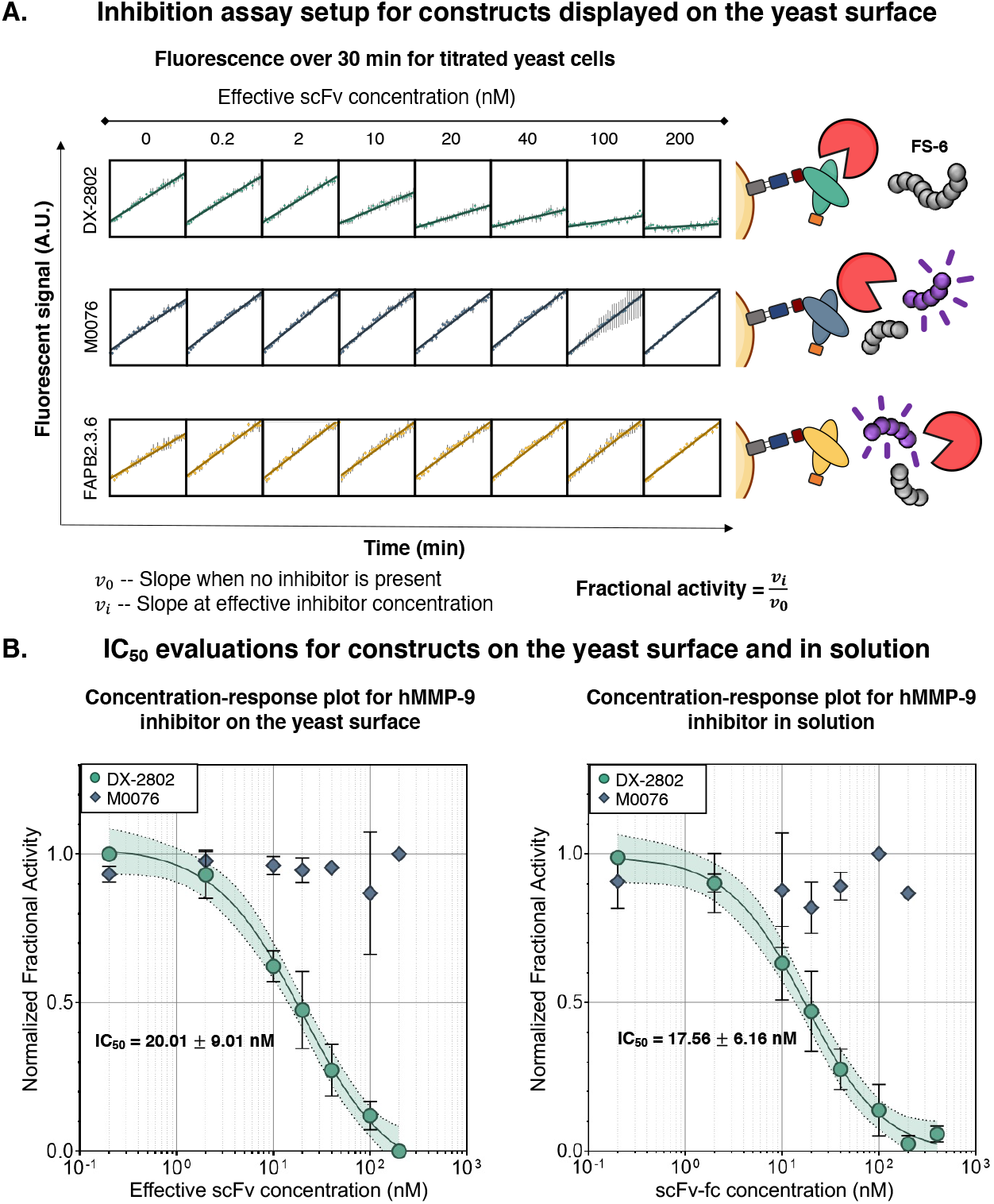
Inhibition assays on the yeast surface and in solution. (**A**) Inhibition assay setup includes titration of scFv-displaying yeast cells. The MMP fluorescent substrate FS-6 fluoresces when cleaved by hMMP-9. Fluorescence over time is obtained for each effective scFv concentration over 30 min and fractional activity for each is obtained from the slopes. (**B**) IC_50_ values for constructs on the yeast surface are calculated from the concentration-response plot (left) using an interpolation curve to fit the data. Concentration-response plot (right) for constructs in solution was obtained from the slope of fluorescence over time.

We used the data in Fig. 5A as the basis for determining the IC_50_ of yeast-displayed DX2802. The rate of increase of fluorescence for each effective scFv concentration and for controls in which no cells were added to wells containing active hMMP-9 enabled determination of fractional enzyme activity as a function of concentration. Following normalization, these data were plotted in concentration-response curves and fitted to a Sigmoidal, 4PL interpolation to determine IC_50_ values (Fig. 5B; see Supplementary Fig. S6 for details). For cells displaying DX-2802, and with a concentration of 2 nM hMMP-9 in solution, we obtained an IC_50_ value of 20.01 ± 9.01 nM, while for cells displaying M0076, the lack of reduction of enzyme activity at all effective concentrations tested precluded fitting.

In order to directly compare these experiments with solution-based measurements, we prepared soluble forms of DX-2802 and M0076 in scFv-Fc format^4^. These two constructs were expressed in secreted form and isolated via protein A affinity chromatography; confirmations of purity were made using SDS-PAGE and Western blotting analyses (see Supplementary Fig. S5 and *Materials and Methods* for details). Titration assays conducted with soluble scFv-Fcs were performed under the same conditions as for the yeast-based titrations, again using 2 nM hMMP-9 in solution (Fig. 5B). Similar to what we observed in yeast-display format, hMMP-9 activity is concentration-dependent only when incubated with increasing amounts of DX-2802. Overall, the experimental errors determined in soluble form are indistinguishable from the errors determined for the data derived from yeast-based titrations. Using the same data analysis as above, we determined an IC_50_ value of 17.56 ± 6.16 nM for DX-2802 in solution (Fig. 5B; see *Materials and Methods* for details). The IC_50_ values obtained in the two different formats fall within the confidence intervals of each other, indicating an excellent level of agreement considering the differences between display and soluble formats (including assumptions of construct display levels). Taken as a whole, these data show that quantitative evaluations of inhibitory properties of clones can be performed directly on the yeast surface.

## Discussion

In this work, we investigated key steps in the discovery and characterization of proteinbased inhibitors using a series of antibody constructs and the therapeutically relevant target MMP-9. The clones used here, MMP-9 inhibitory clone DX-2802, MMP-9 binding clone M0076, and nonbinding control FAPB2.3.6, all exhibited distinct binding and display properties in flow cytometry assays. These varied properties enabled us to rapidly evaluate sorting and characterization approaches. In both bead-based enrichments and fluorescence-activated cell sorting, we were able to preferentially isolate inhibitory clones to hMMP-9 even when binding clones vastly outnumbered inhibitory clones. For bead-based enrichments, we found that preincubation of a population with soluble MMP-9 prior to adding magnetic beads drastically enhanced isolation of inhibitors in comparison to cases where the populations were exposed only to the immobilized target on magnetic beads. Since magnetic bead-based enrichments readily scale to 10 billion cells or more^28^, we expect that this approach could be used even in early-stage enrichments prior to fluorescence-activated cell sorting^25, 26^. FACS-based enrichments resulted in efficient isolation of inhibitory clones with a gating strategy requiring clones to exhibit high cMyc levels in addition to target binding. Starting from model populations with the same initial compositions as used in bead-based procedures, the enrichment of inhibitory binders via FACS was more efficient than enrichment via magnetic bead sorting; this is consistent with previous work in the area^7, 27, 29^. However, the throughput of FACS is typically limited to no more than 10^8^ cells per hour, meaning that the largest practical library size that can be used for FACS enrichments without sacrificing library coverage is 10^7^ unique clones. For yeast display libraries containing more than 10^7^ members, early rounds of enrichments will require the use of bead-based enrichments before more efficient FACS-based enrichments can be used without sacrificing library coverage.

In addition to gaining insights into strategies for enriching for inhibitors, we were also able to obtain IC_50_ values for inhibitors directly on the yeast surface. The values obtained with assays conducted in scFv display format were consistent with the values of assays performed in solution in scFv-Fc format. While previous reports have described titrations of yeast to evaluate inhibition^30^, we are not aware of any work in which IC_50_ values were determined on yeast. Yeast-based assays for IC_50_ value determination are expected to support multiple key evaluations during inhibitor discovery and characterization. First, multiple individual clones could be characterized and rank-ordered based on IC_50_ values in order to quickly prioritize additional characterization efforts, such as which clones should be prepared in soluble format^4, 31^. Second, yeast-based titrations are also expected to support IC_50_ value determinations of libraries or enriched library subpopulations of yeast. This may be useful during library screens to confirm the enrichment of inhibitory clones within a population and even begin to estimate inhibitory potencies at the population level. In principle, yeast-based titrations may support the determination of kinetic parameters of inhibitors, such as the inhibitory constant K_i_. While knowledge of these constants is desirable, since they depend on the mode of inhibition, additional experiments to identify this mode would be necessary prior to determining these constants. On the other hand, the titration-based IC_50_ determinations described here can be performed without knowledge of mode of inhibition, as long as any comparisons between different inhibitors are performed using the same target concentration during IC_50_ value determination (IC_50_ values are a function of the concentration of the target in solution). One important area for future investigation is the extent to which avidity effects influence IC_50_ values determined on the yeast surface. Yeast display is typically performed under conditions that result in high avidity. Such conditions are known to affect quantitative characterizations of binders and inhibitors, particularly when targets are known to be multivalent^32, 33^. Our findings here demonstrate that the evaluation of IC_50_ values directly on the yeast surface enables characterization of target inhibition with data quality that is comparable to data obtained with soluble inhibitor candidates.

Overall, our findings with an MMP-9 model system show that straightforward modifications to enrichment strategies lead to the preferential enrichment of inhibitors, potentially streamlining future high throughput inhibitor discovery. MMP-9 is part of the large matrix metalloproteinase family of proteases, many of which exhibit similar proteolytic capabilities compared to those of MMP-9, as do proteases from several additional families including serine or cysteine proteases^34^. This suggests that our findings could be transferrable to additional targets, although there may be nuances which are target- or application-specific. Extensions to enzymes beyond proteases may also be possible, keeping in mind that specific details of yeast-based enrichments and characterizations may need to be tailored based on the target of interest. Finally, these efforts for isolating protein-based inhibitors in display format could be combined with ongoing efforts to integrate an expanded range of chemical functionality into proteins in search of therapeutic leads^14, 35–41^, or with emerging machine learning approaches to identify protein features that lead to effective protein-based inhibitors^42–44^.

## Acknowledgements

This research was supported by a grant from the National Cancer Institute of the National Institutes of Health (R21CA214239). The content of this work is solely the responsibility of the authors and does not necessarily represent the official views of the National Institutes of Health.

## Materials and Methods

### Plasmid Design and Construction

Sequences encoding single-chain variable fragments (scFvs) of M0076-D03 (in this paper referred to as M0076), a non-inhibitory antibody that binds to human MMP-9 with a reported K_D_ of 5.9 nM^22, 23^, and DX-2802, an antibody that binds to and inhibits human MMP-9^22, 23^ with a reported IC_50_ in the low single-digit nM range (for the specific MMP-9 concentration used during the titration experiment)^45^, were obtained via gene synthesis (GeneArt). Sequences were ordered from GeneArt with an NheI site immediately upstream of the light chain, a BamHI site immediately downstream of the heavy chain, and linker 218 (Supplementary Table S2)^46^ between the light and heavy chain. Sequences were codon-optimized for *S. cerevisiae* by GeneArt. Both sequences were cloned into pCTCON2 for yeast surface display purposes via ligation cloning via the NheI and BamHI sites with pCTCON2 to fuse the scFvs to the C-terminus of Aga2p. The plasmids DNA was chemically transformed into *E. coli* and plated on selective media. Colonies were inoculated into liquid media, grown to saturation, miniprepped and the resulting plasmids were isolated and sequenced. Sequencing for this work was performed by Quintara Biosciences. FAPB2.3.6, an scFv that recognizes both forms of fibroblast activation protein (FAP)^24^, was used as non-binding control in the same yeast display format. Sequences of all display constructs are provided in Supplementary Table S1.

Soluble forms of DX-2802 and M0076 were prepared by cloning the scFv sequences into pCHA-FcSup-TAA (Trp marker) to be secreted as scFv-Fcs and to study their properties in solution^47^. The DX-2802 and M0076 genes were amplified with primers containing the corresponding restriction enzyme sites NheI at the 5’ end and XmaI at the 3’ end, and overlapping regions of the pCHA-FcSup-TAA vector. The vector was digested with the corresponding restriction enzymes, and Gibson assembly was performed to introduce the DX-2802 and M0076 genes into pCHA-FcSup-TAA. The plasmid DNA was transformed into *E. coli* and plated on selective media. Colonies were inoculated into liquid media, grown to saturation, miniprepped and the resulting plasmids were isolated and sequenced. Sequencing for this work was performed by Quintara Biosciences. The resulting plasmids were named pCHA-FcSup-DX-2802 and pCHA-FcSup-M0076.

### Media and Buffer Preparation

The construction of the RJY100 strain of *Saccharomyces cerevisiae*, the host cell for yeast surface display used here, has been described elsewhere^47^. All liquid and solid media were prepared using procedures described in previous work^28^.

MMP buffer (pH7.5, 50mM Tris-HCl, 150mM NaCl, 5mM CaCl_2_), PBSA (pH 7.4, 40 g/L NaCl, 1 g/L KCl, 7.2 g/L Na_2_HPO_4_, 1.2 g/L KH_2_PO_4_, 1 g/L BSA), and PBS (pH 7.4, 40 g/L NaCl, 1 g/L KCl, 7.2 g/L Na_2_HPO_4_, 1.2 g/L KH_2_PO_4_) were prepared and sterile filtered for these experiments.

### MMP-9 Preparation and Buffer Exchange

Freestyle mammalian Human Embryonic Kidney 293 (HEK293F) cells grown in suspension were used to express the human isoform of MMP-9 (henceforth referred to as pro-hMMP-9), using transient DNA transfection. A full length cDNA of human MMP-9 cloned into the pCMV3-C-His vector was used for transfection, resulting in the expression of full length human MMP-9 protease with a C-terminal His-tag (MMP-9 cDNA ORF Clone, Human, C-His tag; SinoBiological cat: HG10327-CH). Transfection-grade plasmid DNA was prepared in milligram quantities by transforming into NEB stable competent *E. coli* (New England Biolabs), growing on solid and liquid LB media supplemented with antibiotic and purifying plasmid DNA using the NucleoBond Xtra Maxi kit (Macherey-Nagel).

To prepare HEK293F cells for transfection, frozen cells were thawed and passaged using FreeStyle™ 293 Expression Medium (Gibco) according to manufacturer’s protocols. Once steady cell growth was established (three to four passages after thawing), cell culture volume was expanded to accommodate large scale production of pro-hMMP-9 by diluting to a cell density of 0.4 × 10^6^ cells/mL with each passage. Approximately 24 hours before transfection, cells were diluted to 0.5 × 10^6^ cells/mL in the final volume for transfection (typically 250 – 300 mL per 1 L flask). On the day of transfection, reagents were pre-warmed to 36.5 °C and prepared as follows. 1 μg DNA was combined with 20 μL OptiPRO™ SFM (Gibco) per mL cell culture and sterile filtered using a 0.2 μM filter. 2 μL 1 mg/mL polyethylenimine (PEI) was combined with 20 μL OptiPRO™ SFM per mL cell culture and sterile filtered using a 0.2 μM filter. The two solutions were incubated separately for 5 minutes, after which they were combined and incubated for an additional 10 minutes to form the final transfection reagent. Once prepared, the transfection reagent was added to the cell culture drop by drop while constantly swirling the flask gently to mix. After transfection, the flasks were returned to the 8.0% CO_2_ incubator at 37 °C, with orbital shaking at 125 RPM, and allowed to grow for seven days during expression and secretion of pro-hMMP-9. All steps involving propagation and transfection of HEK293F cultures were performed in a biosafety cabinet.

After seven days, the cultures were pelleted for 20 minutes at 10,000 rcf in a centrifuge chilled to 4 °C, filtered using a 0.2 μM filter in a biosafety cabinet and adjusted to a pH of 8.0 using a 1:10 dilution of 10× equilibration buffer (500 mM NaH_2_PO_4_, 3 M NaCl, pH 8.0). The filtrate was passed twice through a benchtop low pressure chromatography column containing 1 mL packed high affinity Ni-charged resin (Genscript) per 300 – 500 mL filtered supernatant. Following this, the resin was washed three times with 10 mL 1× equilibration buffer (50 mM NaH_2_PO_4_, 300 mM NaCl, pH 8.0). Pro-hMMP-9 was then eluted using 7 mL 1× elution buffer (50 mM NaH_2_PO_4_, 300 mM NaCl, 250 mM C_3_H_4_N_2_, pH 8.0). The eluant was buffer exchanged to remove the 1× elution buffer using Amicon Ultra-15 centrifugal filter units (30 kDa molecular weight cut-off, Millipore Sigma) into MMP buffer and concentrated. The enzyme concentration was determined using a NanoDrop™ Microvolume UV-Vis Spectrophotometer (Thermo Scientific); purity was evaluated using sodium dodecyl sulfate polyacrylamide gel electrophoresis (SDS-PAGE). For long term storage, pro-hMMP-9 in MMP buffer was mixed with pure glycerol in a 1:1 ratio for a 50% glycerol solution, flash frozen using liquid nitrogen and stored at –80 °C.

Frozen vials of pro-hMMP-9 were rapidly thawed using a room-temperature water bath and buffer exchanged to remove glycerol from the storage solution. Buffer exchange was performed by first equilibrating Amicon Ultra-0.5 centrifugal filter units (30 kDa molecular weight cut-off, Millipore Sigma) with MMP buffer, then centrifuging the enzyme for 10 minutes at 14000 rcf at 4 °C for a total of 6 – 8 spins, each time discarding the flow through and resuspending the enzyme to a final volume of 500 μL in MMP buffer. PBS was used instead of MMP buffer if the enzyme needed to be biotinylated following buffer exchange. The enzyme concentration was determined using a NanoDrop™ Microvolume UV-Vis Spectrophotometer (Thermo Scientific). Pro-hMMP-9 was kept on ice at all times and stored at 4 °C for up to a week.

### MMP-9 Biotinylation

To biotinylate pro-hMMP-9, 100 mM biotin solution was prepared by dissolving an appropriate amount of NHS-LC EZ Link Biotin powder (Thermo Scientific) into a volume of dimethylformamide (Sigma Aldrich) at the ratio of 1.5 mg NHS-Lc EZ Link Biotin powder to 32.61 μL of DMF. 100 μL of pro-hMMP-9 protein stock in PBS at 1 mg/mL was incubated with 5–6 fold molar excess of 100 mM biotin solution, mixed well, and incubated on ice for 90 minutes with mixing every 20–30 minutes. The reaction was terminated by adding 10 μL Tris quench buffer (0.5 M Tris, 0.02 % NaN_3_, pH 7.4), mixing well, and letting the reaction sit on ice for 10 minutes. Zeba Spin Desalting Columns with 7K MW cutoff (Thermo Fisher) were equilibrated with PBS and used to remove unreacted biotin after the reaction following the manufacturer’s protocols.

Following protein desalting, the extent of biotinylation was determined using HABA assays. HABA/Avidin premix (Thermo Scientific) was reconstituted in sterile water and 20 μL was added to a clear flat bottom 96-well plate, with 160 μL of PBS buffer. After mixing, endpoint absorbance at 500 nm was measured using a SpectraMax i3x Multi-Mode Microplate Reader (Molecular Devices) until the values were determined to be stable (at least two measurements within ±0.003 AU). 20 μL biotinylated pro-hMMP-9 was added to the mixture, mixed, and absorbance at 500 nm was measured again until the values were determined to be stable. The extent of biotinylation was calculated using the online HABA Calculator. Samples with biotin to protein ratios of 1 to 2 were used immediately or stored at 4 °C for no more than one week.

### MMP-9 Activation and Deactivation

To activate pro-hMMP-9, 0.5 mg/mL Trypsin (Sigma Aldrich) solution in MMP buffer and 100 mM phenylmethylsulfonyl fluoride (PMSF, Sigma Aldrich) solution in isopropanol were thawed and kept on ice until use. Buffer exchanged pro-hMMP-9 (1 mg/mL), Trypsin (0.5 mg/mL) and MMP buffer were incubated at room temperature for 25 minutes at a volumetric ratio of 1:2:7 in 1 mL total volume. Following incubation, 100 mM PMSF at a volumetric ratio of 1:1 with hMMP-9 was added to terminate the reaction using a 5-min incubation period. The enzyme was diluted in the appropriate buffer (200 nM for binding experiments, 2 nM for inhibition assays on the yeast surface) and kept on ice until ready to be used. To validate the activation, 50× solution of the MMP fluorescent substrate FS-6 (Sigma Aldrich, 20 mM in TCNB: 50 mM Tris-Cl, 10 mM CaCl_2_ dihydrate, 150 mM NaCl, 0.5% Brij-35) was thawed and kept on ice and in the dark until ready to be used. Activation was validated by incubating 10 μL of 200 nM activated MMP-9 with 50 μL of 250 μM FS-6, in a clear flat bottom 96-well plate. Fluorescence was measured for a blank control (buffer only), pro-hMMP-9 control, and activated hMMP-9 using the SpectraMax i3x Multi-Mode Microplate Reader (Molecular Devices) with excitation at 328nm and emission at 393nm approximately every 10 s for a total of 10 min.

To obtain a 200 nM solution of deactivated hMMP-9 (deact-hMMP-9) for binding assays, 250 nM activity-verified hMMP-9 was incubated with 0.5M EDTA (final 0.1M) at a volumetric ratio of 4:1 for 30 min at room temperature. Deactivation was verified with a microplate reader as above.

### Propagation and Induction

The plasmids pCTCON2-DX-2802, pCTCON2-M0076, and pCTCON2-FAPB2.3.6 were transformed into competent RJY100 cells, plated on SD-SCAA agar plates, and left to grow at 30 °C for three days. Single colonies were inoculated into SD-CAA liquid media and allowed to grow at 30 °C with shaking to saturation (2-3 days). The cultures were then diluted to an OD_600_ of 1 in fresh SD-CAA and grown at 30 °C until mid-log phase (OD_600_ of 2–5; approximately 4–8 hours). Cells were removed, pelleted (spun for 30 s at 14,000 rcf if in 1.7 mL tubes or 5 min at 2,400 rcf in V-bottom plates) and resuspended to an OD_600_ of 1 in SG-CAA induction media (for example: 2 × 10^6^ cells in 2 mL induction media). All samples were induced at 20 °C for 16 hours with shaking.

### Flow Cytometry

Cell cultures were propagated and induced as described above. 2 × 10^6^ induced cells per sample were transferred to microcentrifuge tubes and pelleted for 30 s at 14,000 rcf and 4 °C. For active hMMP-9 (act-hMMP-9) binding evaluation, samples were washed three times in 1 mL MMP buffer and resuspended in 50 μL of 200nM act-hMMP-9 in MMP buffer. The samples were incubated for 10 min at room temperature on a rotary wheel at 150 RPM. Following this incubation, the samples were washed three times in 1 mL ice-cold PBSA and cells were left as pellets on ice. Primary antibody labelling was prepared with a 1:500 dilution of chicken anti-cMyc and mouse anti-His in PBSA (Supplementary Table S2). The cells were resuspended and incubated with 50 μL of the primary antibody labelling solution at room temperature on a rotary wheel at 150 RPM for 30 min^48^. Cells were then pelleted, washed twice in 1 mL ice-cold PBSA, and were left as pellets on ice. Secondary antibody labelling was prepared with a 1:500 dilution of fluorophores Alexa 488 anti-mouse and Alexa 647 anti-chicken in ice-cold PBSA (Supplementary Table S2). Cells were resuspended and incubated with 50 μL of the secondary antibody labelling solution and left on ice in the dark for 15 min. Following secondary labelling, the cells were washed once with ice-cold PBSA and left as pellets on ice. Immediately prior to running each sample on the flow cytometer, the cells were resuspended in 1 mL ice-cold PBSA. Flow cytometry was performed on an Attune NxT flow cytometer in the Tufts University Science and Technology Center. Results from flow cytometry experiments were analyzed using FlowJo v10.6.2 software.

### Bead-based Enrichments

To prepare model mixtures of cells displaying DX-2802, M0076, and FAPB2.3.6 (ratios of 1:1:100, 1:1:1000, 1:100:100, 1:1000:1000), cells containing plasmids encoding each construct were separately inoculated, grown and induced as described above. Following induction, each culture was pelleted, washed 3× with PBSA and the OD_600_ of the resuspended cells was measured. Volumes of cells corresponding to the total cell numbers needed to generate the four compositions listed above to generate a total of 3 × 10^8^ cells per composition were added to a single tube to account for triplicates. The resulting sets of cells were mixed well by vortexing prior to splitting each sample into separate tubes containing 10^8^ cells per tube. Cells were then pelleted and placed on ice.

#### Bead preparation and enrichment with antigen coated beads

Biotin Binder DynaBeads® (Thermo Fisher) were washed and prepared as described elsewhere^28^. Briefly, 10 μL of biotin binder beads per sample were incubated with 33 pmol biotinylated pro-hMMP-9 and 100 μL PBSA in a 1.7 mL Eppendorf tube for 2 hours at 4 °C on a rotary wheel. At the end of the incubation period, the magnetic beads were washed twice by adding 1 mL PBSA, placing the tube on a Dynamag-2 magnet for 2 min, and removing the supernatant^28^ to remove any unbound antigen. After the washes, the beads were resuspended in the original volume of PBSA and kept on ice. The pro form of hMMP-9 coated on beads was activated using Trypsin as described above. Following activation with Trypsin and incubation with PMSF, the beads were washed twice as above on a Dynamag-2 magnet for 2 min and resuspended in the original volume of PBSA. The activity of the resulting immobilized hMMP-9 was determined by incubating the immobilized enzyme with the fluorescent substrate (FS-6)^49^, and fluorescence was measured once approximately every 10 s for a total of 20 minutes on the plate reader as described above. Following activation, 10 μL act-hMMP-9-coated beads were incubated with the washed cell mixtures in 1 mL PBSA for 30 min at RT on a rotary wheel. Immediately following incubation, beads were washed twice in PBSA on a magnet^28^ and rescued in selective growth media (SD-CAA). After growing the recovered beads (and bound cells) overnight at 30 °C with shaking at 300 RPM, the magnetic beads were removed and the cultures propagated, diluted, and induced to prepare for flow cytometry characterization as described above.

#### Bead preparation and enrichment with antigen in solution

Biotin Binder DynaBeads® (Thermo Fisher) were washed and prepared as described elsewhere^28^. Pro-hMMP-9 in solution was activated using Trypsin as described above. The activation was validated using FS-6 as described above. Washed cells were incubated with 200nM act-hMMP-9 for 30 minutes at RT on a rotary wheel.

Immediately following incubation, samples were washed 3 times with PBSA (30 seconds at 14000 rcf) to remove unbound MMP, resuspended in 1 mL of PBSA and incubated with 10 μL biotin-binder beads per sample for 2 hours at 4 °C on a rotary wheel. Following incubation, the beads were washed twice on a Dynamag-2 magnet as described above and rescued in SDCAA. After growing the recovered beads (and bound cells) overnight at 30 °C with shaking at 300 RPM, the magnetic beads were removed and the cultures propagated, diluted, and induced to prepare for flow cytometry characterization as described above.

To isolate plasmids encoding individual clones from enriched samples, cells from each rescued culture (ratio 1:1:100, 1:1:1000, 1:100:100, 1:1000:1000 enriched via bead enrichment with antigen in solution) were grown and yeast miniprepped using Zymoprep™ yeast miniprep protocols. The plasmids obtained were transformed into competent *E. coli* DH5αZ1 strain and plated on LB+Amp plates. 10 colonies from each plate were grown in liquid media, miniprepped and plasmids were sent for sequencing at Quintara Biosciences.

### Fluorescence Activated Cell Sorting

To prepare the two compositions for enrichments using fluorescence activated cell sorting (FACS) using cells displaying DX-2802, M0076, or FAPB2.3.6 (ratios of 1:100:100 and 1:1000:1000), each culture was separately inoculated, grown and induced for 20 hours as described above. Following induction, each culture was pelleted, washed 3× with PBSA and the OD_600_ of the resuspended cells was measured. Cells were mixed corresponding to the two ratios, with each composition having 3 × 10^7^ cells to account for technical triplicates. Prior to incubation and labelling, each composition was divided into three replicates with each sample having 10^7^ cells. Each sample was incubated with 20 μL of 200 nM act-hMMP-9 at room temperature for 10 min on a rotary wheel at 150 RPM. Following incubation, cells were washed 3× with MMP buffer and 3× with EDTA to remove unbound hMMP-9 and stop any further MMP activity. Samples were separately labelled for flow cytometry as described above, and populations were separated via FACS based on relative levels of cMyc and hMMP-9 detection using a BD Aria II FACS machine (BD Biosciences) at the Tufts University Flow Cytometry Core. Following sorting, the cells were rescued in SDCAA at 30 °C with shaking at 300 RPM. Upon saturation, cultures were diluted and induced to prepare for flow cytometry characterization as described above.

To isolate plasmids encoding individual clones from enriched samples, cells from each rescued culture (ratio 1:100:100, 1:1000:1000 enriched via FACS) were grown and yeast miniprepped. The plasmids obtained were transformed into competent *E. coli* DH5αZ1, plated on LB+Amp plates, and allowed to grow at 37 °C. 10 colonies from each plate were grown in liquid media, miniprepped, and plasmids were sent for sequencing at Quintara Biosciences.

### Titrations with Yeast Displaying Antibodies

Cells transformed with the plasmids encoding for yeast displayed versions of DX-2802, M0076, and FAPB2.3.6 were inoculated, diluted, and induced as described above. Induced cells were washed 3× in MMP buffer and the OD_600_ of the resuspended cells was measured following washing. Making the assumption that each yeast cell displays 5×10^4^ constructs following induction^6, 28^, cells displaying the constructs were added to triplicate sets of wells of a V-bottom 96-well plate at increasing amounts. The number of cells chosen for each effective concentration was determined based on the concentration ratio between displayed constructs and hMMP-9. Supplementary Table S3 shows the number of cells added in each well of the first column of a 96-well plate, and the corresponding effective concentration and concentration ratio with hMMP-9. Each set was run in technical triplicates in the same plate. The following equation was used to calculate the effective concentration:

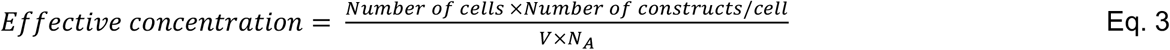

where V is the sample volume and N_A_ is Avogadro’s number. Displaying cells were then pelleted and resuspended in 100 μL of 2 nM active hMMP-9 in MMP buffer and incubated for 10 minutes in a shaking platform at 150 RPM. Immediately following incubation, 20 μL of each sample was taken from the V-bottom 96-well plate and placed into a black-well Costar flat clear-bottom 96-well plate. Each sample was mixed with 100 μL of 50 μM MMP fluorescent substrate (FS-6) and fluorescence was measured once approximately every 10 s for a total of 20 minutes on the SpectraMax i3x Multi-Mode Microplate Reader (Molecular Devices) with excitation at 328 nm and emission at 393 nm with 10 s orbital shaking within each read. Each experiment was run in technical triplicates where cells were grown and induced separately, and the experiment was conducted on different days.

Fluorescence over time for each sample was measured on the spectrophotometer and the values were exported and analyzed. For each well, the slope of fluorescence vs. time was determined. For each well that contained yeast cells (rows B-H) the slope (*v_i_*) was compared to the slope (*v*_0_) of the MMP only control (row A) to obtain fractional activity following Equation 4.

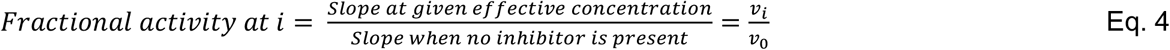

The data from each sample was then analyzed in two ways: (1) normalized in Excel through Equation 5, then interpolated via GraphPad Prism v9.1 using Sigmoidal, 4PL parameters, and (2) normalized in GraphPad Prism v9.1, then interpolated using Sigmoidal, 4PL parameters to obtain IC_50_ values.

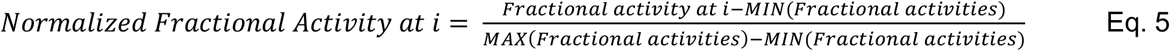

Both methods of analysis yielded matching results. An example of the obtained data and subsequent analysis is described in Supplementary Fig. S6.

### Preparation of soluble constructs

The designed secretion plasmids pCHA-FcSup-DX-2802 and pCHA-FcSup-M0076 described in *Plasmid Design and Construction* were transformed into Zymo-competent RJY100 cells, plated on selective media (–Trp –Ura), and grown at 30 °C until colonies appeared (~3 days). Colonies were picked from each plate and inoculated into 5 mL liquid cultures, which were grown to saturation. The saturated cultures were passaged once, diluted into 50 mL of minimal media, and induced in 100 mL YPG + Pen/Strep for 4 days at 20 °C with shaking at 300 RPM. At the end of induction, the cultures were pelleted for 35 minutes at 3214 rcf and filtered using a 0.2 μm filter. The filtrate was run through a Protein A column twice, and the scFv-Fc containing resin was washed three times with 10 mL 1× PBS pH7.0. The scFv-Fc was eluted in 7 mL 100 mM Glycine pH 3.0 and immediately neutralized with 7 mL 1 M Tris pH 8.5. The eluant was buffer exchanged into 1× PBS pH 7.4 using Amicon Ultra-15 centrifugal filter units (30 kDa MW cut-off, Millipore Sigma) and concentrated. Protein concentrations were measured via 280 nm light absorbance on a NanoDrop One instrument (Thermo Fisher). SDS-PAGE was run to confirm the molecular weight of the purified proteins. A duplicate SDS-PAGE gel was run and the proteins on the gel were transferred onto a nitrocellulose membrane using iBlot™ 2 Gel Transfer Device and iBlot™ transfer stacks according to the manufacturer’s protocols. The membrane was then placed in 5% milk and TBS-T blocking solution and incubated at room temperature with shaking at 60 RPM for 1 h. Following blocking, the membrane was washed twice for 5 minutes each at room temperature with shaking at 60 RPM in enough TBS-T to cover it. The membrane was incubated with PE anti-HA antibody in 1:1000 dilution in 20 mL TBS-T for 1 h at 4 °C with shaking at 60 RPM. The membrane was washed three times for 10 minutes each in TBS-T and was visualized using an Azure c400 gel imager with excitation wavelength 488 nm. SDS-PAGE and Western Blots can be seen in Supplementary Fig. S6. All soluble proteins were mixed in a 1:1 volumetric ratio with glycerol, aliquoted in 200 μL volumes, flash frozen with liquid nitrogen, and stored at – 80 °C for long-term storage.

### Titration in solution

Purified proteins were titrated at eight different concentrations in wells on a V-bottom 96-well plate in triplicate with 50 μL per well at twice the intended final concentration. 50 μL of 4 nM active hMMP-9 in MMP buffer was added to each protein-containing well for a final concentration of 2 nM hMMP-9. The final scFv-Fc concentrations matched the effective concentration of proteins in yeast display format (Supplementary Table S3). The mixture was incubated for 10 minutes in a shaking platform at 150 RPM. Immediately following incubation, 20 μL of each sample was taken from the V-bottom 96-well plate and placed into a black-well Costar flat clear-bottom 96-well plate. Each sample was mixed with 100 μL of 50 μM FS-6 and fluorescence was measured once approximately every 10 s for a total of 20 minutes on the SpectraMax i3x Multi-Mode Microplate Reader (Molecular Devices) with excitation at 328nm and emission at 393nm with 10s orbital shaking within each read. Fractional activity was analyzed as before (see *Titrations with Yeast Displaying Antibodies*), using Equation 4, and normalization was performed in two ways as before, using Equation 5. Visualization and statistical comparisons of IC_50_ values between yeast display constructs and soluble constructs were performed using GraphPad Prism v9.1.

## References

1. Bradbury, A. R.; Sidhu, S.; Dubel, S.; McCafferty, J., Beyond natural antibodies: the power of in vitro display technologies. Nat Biotechnol 2011, 29 (3), 245–54.

2. Bowers, P. M.; Horlick, R. A.; Neben, T. Y.; Toobian, R. M.; Tomlinson, G. L.; Dalton, J. L.; Jones, H. A.; Chen, A.; Altobell, L., 3rd; Zhang, X.; Macomber, J. L.; Krapf, I. P.; Wu, B. F.; McConnell, A.; Chau, B.; Holland, T.; Berkebile, A. D.; Neben, S. S.; Boyle, W. J.; King, D. J., Coupling mammalian cell surface display with somatic hypermutation for the discovery and maturation of human antibodies. Proc Natl Acad Sci U S A 2011, 108 (51), 20455–60.

3. Cherf, G. M. C. J. R., Applications of Yeast Surface Display for Protein Engineering. Yeast Surface Display: Methods, Protocols, and Applications. 2015, (1319), 20.

4. Van Deventer, J. A.; Le, D. N.; Zhao, J.; Kehoe, H. P.; Kelly, R. L., A platform for constructing, evaluating, and screening bioconjugates on the yeast surface. Protein Eng Des Sel 2016, 29 (11), 485–494.

5. Konning, D.; Kolmar, H., Beyond antibody engineering: directed evolution of alternative binding scaffolds and enzymes using yeast surface display. Microb Cell Fact 2018, 17 (1), 32.

6. Chao, G.; Lau, W. L.; Hackel, B. J.; Sazinsky, S. L.; Lippow, S. M.; Wittrup, K. D., Isolating and engineering human antibodies using yeast surface display. Nat Protoc 2006, 1 (2), 755–68.

7. Chen, T. F.; de Picciotto, S.; Hackel, B. J.; Wittrup, K. D., Engineering fibronectin-based binding proteins by yeast surface display. Methods Enzymol 2013, 523, 303–26.

8. Bacon, K.; Blain, A.; Bowen, J.; Burroughs, M.; McArthur, N.; Menegatti, S.; Rao, B. M., Quantitative Yeast-Yeast Two Hybrid for the Discovery and Binding Affinity Estimation of Protein-Protein Interactions. ACS Synth Biol 2021, 10 (3), 505–514.

9. Devaraj, N. K., The Future of Bioorthogonal Chemistry. ACS Cent Sci 2018, 4 (8), 952–959.

10. Denard, C. A.; Paresi, C.; Yaghi, R.; McGinnis, N.; Bennett, Z.; Yi, L.; Georgiou, G.; Iverson, B. L., YESS 2.0, a Tunable Platform for Enzyme Evolution, Yields Highly Active TEV Protease Variants. ACS Synth Biol 2021, 10 (1), 63–71.

11. Jordan, K. R.; McMahan, R. H.; Oh, J. Z.; Pipeling, M. R.; Pardoll, D. M.; Kedl, R. M.; Kappler, J. W.; Slansky, J. E., Baculovirus-infected insect cells expressing peptide-MHC complexes elicit protective antitumor immunity. J Immunol 2008, 180 (1), 188–97.

12. Galan, A.; Comor, L.; Horvatic, A.; Kules, J.; Guillemin, N.; Mrljak, V.; Bhide, M., Library-based display technologies: where do we stand? Mol Biosyst 2016, 12 (8), 2342–58.

13. Boder, E. T.; Raeeszadeh-Sarmazdeh, M.; Price, J. V., Engineering antibodies by yeast display. Arch Biochem Biophys 2012, 526 (2), 99–106.

14. Islam, M.; Kehoe, H. P.; Lissoos, J. B.; Huang, M.; Ghadban, C. E.; Berumen Sanchez, G.; Lane, H. Z.; Van Deventer, J. A., Chemical Diversification of Simple Synthetic Antibodies. ACS Chem Biol 2021, 16 (2), 344–359.

15. Arkadash, V.; Yosef, G.; Shirian, J.; Cohen, I.; Horev, Y.; Grossman, M.; Sagi, I.; Radisky, E. S.; Shifman, J. M.; Papo, N., Development of High Affinity and High Specificity Inhibitors of Matrix Metalloproteinase 14 through Computational Design and Directed Evolution. J Biol Chem 2017, 292 (8), 3481–3495.

16. Christmann, A.; Walter, K.; Wentzel, A.; Kratzner, R.; Kolmar, H., The cystine knot of a squash-type protease inhibitor as a structural scaffold for *Eschirichia coli* cell surface display of conformationally constrained peptides. Protein Engineering 1999, 12 (9), 9.

17. Lee, K. B.; Nam, D. H.; Nuhn, J. A. M.; Wang, J.; Schneider, I. C.; Ge, X., Direct expression of active human tissue inhibitors of metalloproteinases by periplasmic secretion in Escherichia coli. Microb Cell Fact 2017, 16 (1), 73.

18. Mehta, N.; Maddineni, S.; Kelly, R. L.; Lee, R. B.; Hunter, S. A.; Silberstein, J. L.; Parra Sperberg, R. A.; Miller, C. L.; Rabe, A.; Labanieh, L.; Cochran, J. R., An engineered antibody binds a distinct epitope and is a potent inhibitor of murine and human VISTA. Sci Rep 2020, 10 (1), 15171.

19. Papo, N.; Silverman, A. P.; Lahti, J. L.; Cochran, J. R., Antagonistic VEGF variants engineered to simultaneously bind to and inhibit VEGFR2 and alphavbeta3 integrin. Proc Natl Acad Sci U S A 2011, 108 (34), 14067–72.

20. Raeeszadeh-Sarmazdeh, M.; Coban, M.; Mahajan, S.; Hockla, A.; Sankaran, B.; Downey, G. P.; Radisky, D. C.; Radisky, E. S., Engineering of tissue inhibitor of metalloproteinases TIMP-1 for fine discrimination between closely related stromelysins MMP-3 and MMP-10. Journal of Biological Chemistry 2022, 298 (3), 101654.

21. Frenzel, A.; Schirrmann, T.; Hust, M., Phage display-derived human antibodies in clinical development and therapy. MAbs 2016, 8 (7), 1177–1194.

22. Wood, C. R. Combination treatments comprising protease binding proteins for inflammatory disorders. 2011.

23. Nicholson, S.; Wood, C.; Devy, L. Use of MMP-9 and MMP-12 binding proteins for the treatment and prevention of systemic sclerosis. 2010.

24. Stieglitz, J. T.; Kehoe, H. P.; Lei, M.; Van Deventer, J. A., A Robust and Quantitative Reporter System To Evaluate Noncanonical Amino Acid Incorporation in Yeast. ACS Synth Biol 2018, 7 (9), 2256–2269.

25. Angelini, A.; Chen, T. F.; de Picciotto, S.; Yang, N. J.; Tzeng, A.; Santos, M. S.; Van Deventer, J. A.; Traxlmayr, M. W.; Wittrup, K. D., Protein Engineering and Selection Using Yeast Surface Display. In Yeast Surface Display: Methods, Protocols, and Applications, Liu, B., Ed. Springer New York: New York, NY, 2015; pp 3–36.

26. Gera, N.; Hussain, M.; Rao, B. M., Protein selection using yeast surface display. Methods 2013, 60 (1), 15–26.

27. Volden, T. A.; Reyelts, C. D.; Hoke, T. A.; Arikkath, J.; Bonasera, S. J., Validation of Flow Cytometry and Magnetic Bead-Based Methods to Enrich CNS Single Cell Suspensions for Quiescent Microglia. Journal of Neuroimmune Pharmacology 2015, 10 (4), 655–665.

28. Van Deventer, J. A.; Wittrup, K. D., Yeast surface display for antibody isolation: library construction, library screening, and affinity maturation. Methods Mol Biol 2014, 1131, 151–81.

29. Colby, D. W.; Kellogg, B. A.; Graff, C. P.; Yeung, Y. A.; Swers, J. S.; Wittrup, K. D., Engineering Antibody Affinity by Yeast Surface Display. In Methods in Enzymology, Academic Press: 2004; Vol. 388, pp 348–358.

30. Bowen, J.; Schneible, J.; Bacon, K.; Labar, C.; Menegatti, S.; Rao, B. M., Screening of Yeast Display Libraries of Enzymatically Treated Peptides to Discover Macrocyclic Peptide Ligands. Int J Mol Sci 2021, 22 (4).

31. Cruz-Teran, C. A.; Bacon, K.; Rao, B. M., Simultaneous Soluble Secretion and Surface Display of Proteins in Saccharomyces cerevisiae Using Inefficient Ribosomal Skipping. Methods Mol Biol 2020, 2070, 321–334.

32. Dubacheva, G. V.; Araya-Callis, C.; Geert Volbeda, A.; Fairhead, M.; Codée, J.; Howarth, M.; Richter, R. P., Controlling Multivalent Binding through Surface Chemistry: Model Study on Streptavidin. Journal of the American Chemical Society 2017, 139 (11), 4157–4167.

33. Lacham-Hartman, S.; Shmidov, Y.; Radisky, E. S.; Bitton, R.; Lukatsky, D. B.; Papo, N., Avidity observed between a bivalent inhibitor and an enzyme monomer with a single active site. PLOS ONE 2021, 16 (11), e0249616.

34. Patel, S., A critical review on serine protease: Key immune manipulator and pathology mediator. Allergol Immunopathol (Madr) 2017, 45 (6), 579–591.

35. Lewis, A. K.; Harthorn, A.; Johnson, S. M.; Lobb, R. R.; Hackel, B. J., Engineered protein-small molecule conjugates empower selective enzyme inhibition. Cell Chem Biol 2021.

36. Passioura, T.; Liu, W.; Dunkelmann, D.; Higuchi, T.; Suga, H., Display Selection of Exotic Macrocyclic Peptides Expressed under a Radically Reprogrammed 23 Amino Acid Genetic Code. J Am Chem Soc 2018, 140 (37), 11551–11555.

37. Rezhdo, A.; Islam, M.; Huang, M.; Van Deventer, J. A., Future prospects for noncanonical amino acids in biological therapeutics. Curr Opin Biotechnol 2019, 60, 168–178.

38. Schellenberger, V.; Wang, C. W.; Geething, N. C.; Spink, B. J.; Campbell, A.; To, W.; Scholle, M. D.; Yin, Y.; Yao, Y.; Bogin, O.; Cleland, J. L.; Silverman, J.; Stemmer, W. P., A recombinant polypeptide extends the in vivo half-life of peptides and proteins in a tunable manner. Nat Biotechnol 2009, 27 (12), 1186–90.

39. Navaratna, T.; Atangcho, L.; Mahajan, M.; Subramanian, V.; Case, M.; Min, A.; Tresnak, D.; Thurber, G. M., Directed Evolution Using Stabilized Bacterial Peptide Display. Journal of the American Chemical Society 2020, 142 (4), 1882–1894.

40. Babin, B. M.; Keller, L. J.; Pinto, Y.; Li, V. L.; Eneim, A. S.; Vance, S. E.; Terrell, S. M.; Bhatt, A. S.; Long, J. Z.; Bogyo, M., Identification of covalent inhibitors that disrupt M. tuberculosis growth by targeting multiple serine hydrolases involved in lipid metabolism. Cell Chemical Biology 2021.

41. Ekanayake, A. I.; Sobze, L.; Kelich, P.; Youk, J.; Bennett, N. J.; Mukherjee, R.; Bhardwaj, A.; Wuest, F.; Vukovic, L.; Derda, R., Genetically Encoded Fragment-Based Discovery from Phage-Displayed Macrocyclic Libraries with Genetically Encoded Unnatural Pharmacophores. Journal of the American Chemical Society 2021, 143 (14), 5497–5507.

42. Hassoun, S.; Jefferson, F.; Shi, X.; Stucky, B.; Wang, J.; Rosa, E., Artificial Intelligence for Biology. Integr Comp Biol 2021.

43. Li, X.; Van Deventer, J.; Hassoun, S., Towards the Design of Matrix Metalloproteinases (MMP) Antibody Sequences. In Proceedings of the 8th ACM International Conference on Bioinformatics, Computational Biology,and Health Informatics, 2017; pp 624–624.

44. Mahajan, S. P.; Meksiriporn, B.; Waraho-Zhmayev, D.; Weyant, K. B.; Kocer, I.; Butler, D. C.; Messer, A.; Escobedo, F. A.; DeLisa, M. P., Computational affinity maturation of camelid single-domain intrabodies against the nonamyloid component of alpha-synuclein. Sci Rep 2018, 8 (1), 17611.

45. Beck, A.; Reichert, J. M.; Wurch, T., 5th European Antibody Congress 2009: November 30-December 2, 2009, Geneva, Switzerland. MAbs 2010, 2 (2), 108–28.

46. Tran, E.; Chinnasamy, D.; Yu, Z.; Morgan, R. A.; Lee, C.-C. R.; Restifo, N. P.; Rosenberg, S. A., Immune targeting of fibroblast activation protein triggers recognition of multipotent bone marrow stromal cells and cachexia. Journal of Experimental Medicine 2013, 210 (6), 1125–1135.

47. Van Deventer, J. A.; Kelly, R. L.; Rajan, S.; Wittrup, K. D.; Sidhu, S. S., A switchable yeast display/secretion system. Protein Engineering, Design and Selection 2015, 28 (10), 317–325.

48. Stieglitz, J. T. V. D., J. A., Incorporating, quantifying, and leveraging noncanonical amino acids in yeast. Methods Mol Biol Accepted.

49. Neumann, U.; Kubota, H.; Frei, K.; Ganu, V.; Leppert, D., Characterization of Mca-Lys-Pro-Leu-Gly-Leu-Dpa-Ala-Arg-NH2, a fluorogenic substrate with increased specificity constants for collagenases and tumor necrosis factor converting enzyme. Anal Biochem 2004, 328 (2), 166–73.

