## Supplementary Information for "Strategies for enriching and characterizing proteins with inhibitory properties on the yeast surface"

**Authors/Affiliations**

Arlinda Rezhdo[†], Catherine T. Lessard[†], Mariha Islam[†], and James A. Van Deventer[*, †, ‡]

[†]Chemical and Biological Engineering Department, Tufts University, Medford, Massachusetts 02155, USA

[‡]Biomedical Engineering Department, Tufts University, Medford, Massachusetts 02155, USA

*Corresponding author
;

### Supplementary Tables

**Table S1.** Sequence information for the three constructs used in this paper from the beginning of the NheI restriction site (highlighted in yellow) to the end of the BamHI restriction site (highlighted in blue) with linker 218 (highlighted in gray).

| scFv construct | Sequence |
|---|---|
| **DX-2802** | GCTAGCCAATCTGCTTTGACTCAACCTAGATCTGTTTCTGGTTCTCCAGGTCAA<br>TCTGTTACTATTTCTTGTACTGGTACTTCCTCTGATGTTGGTGGTTACAATTACG<br>TCAGTTGGTATCAACAACATCCAGGTAAAGCTCCAAAGTTGATGATCTACGATG<br>TCTCCAAAAGACCATCTGGTGTTCCAGATAGATTCTCTGGTTCTAAGTCTGGTA<br>ATACTGCCTCTTTGACCATCTCTGGTTTACAAGCTGAAGATGAAGCTGATTACT<br>ACTGTTGTTCTTACGCTGGTTCTTACACTTTGGTTTTTGGTGGTGGTACTAAGT<br>TGACTGTTTTGGGCTCCACTAGCGGTTCCGGCAAACCTGGCAGCGGAGAAGG<br>CAGCACCAAAGGGGAAGTTCAATTATTGGAATCAGGTGGTGGTTTGGTTCAAC<br>CAGGTGGTTCTTTGAGATTGTCTTGTGCTGCTTCTGGTTTCACTTTCTCTACTT<br>ATCAAATGGTCTGGGTTAGACAAGCTCCTGGTAAAGGTTTGGAATGGGTTTCT<br>GTTATCTATCCTTCTGGTGGTCCAACTGTTTATGCTGATTCTGTTAAGGGTAGA<br>TTCACCATCTCCAGAGATAATTCTAAGAACACCTTGTACTTGCAAATGAACTCC<br>TTGAGAGCAGAAGATACTGCTGTTTATTACTGTGCTAGAGGTGAAGATTACTAC<br>GATTCTTCTGGTCCAGGTGCTTTTGATATTTGGGGTCAAGGTACTATGGTTACC<br>GTTTCTTCTGGATCC |
| **M0076** | GCTAGCCAAGATATTCAAATGACCCAATCTCCTTCTTCCTTGTCTGCTTCTGTT<br>GGTGATAGAGTTACTATTACCTGTAGAGCCTCTCAAGGTATCAGAAATGATTTG<br>GACTGGTATCAACAAAAGCCAGGTACTGCTCCAAAAAGATTGATCTATTCTGCC<br>TCCAACTTGCAATCTGGTGTTCCATCTAGATTTTCTGGTTCTGGTTCAGGTACT<br>GAATTTACCTTGACCATCTCTAACTTGCAACCAGAAGATTTGGCTACTTACTTC<br>TGCTTGCAACATAACTCTTTCCCATTGACTTTCGGTCAAGGTACTAAGGTTGAA<br>ATCAAAGGCTCCACTAGCGGTTCCGGCAAACCTGGCAGCGGAGAAGGCAGCA<br>CCAAAGGGGAAGTTCAATTATTGGAATCTGGTGGTGGTTTGGTTCAACCAGGT<br>GGTTCTTTGAGATTGTCTTGTGCTGCTTCAGGTTTCACCTTCTCATTATACAGA<br>ATGAACTGGGTTAGACAAGCTCCAGGTAAAGGTTTGGAATGGGTTTCTTATATT<br>GGTTCTTCAGGTGGTGCTACTGCTTATGCTGATTCTGTTAAGGGTAGATTCACC<br>ATCTCCAGAGATAATTCTAAGAACACCTTGTACTTGCAAATGAACTCCTTGAGA<br>GCTGAAGATACTGCTGTTTATTACTGTGCTAGAGGTGCTTGGTATTTGGATTCT<br>TGGGGTCAAGGTACTTTGGTTACTGTTTCTTCTGGATCC |
| **FAPB2.3.6** | GCTAGCGATATCCAGATGACCCAGTCCCCGAGCTCCCTGTCCGCCTCTGTGA<br>GCGATAGGGTCACCATCACCTGCCGTGCCGGTCAGTACGGTTCTTCTTACGTA<br>GCCTGGTATCAACAGAAACCAGGAAAAGCTCCGAAGCTTCTGATTTACTCTGC<br>ATCCCACCTCTACTCTGGAGTCCCTTCTCGCTTCTCTGGTAGCCGTTCCGGGA<br>CGGGTTTCACTCTGACCATCAGCAGTCTGCAGCCGGAAGACTTCGCAACTTAT<br>TACTGTCAGCAACCGGGTCCGGCTCCGGGTCTGATCACGTTCGGACAGGGTA<br>CCAAGGTGGAGATCAAAGGTACCACTGCCGCTAGTGGTAGTAGTGGTGGCAG<br>TAGCAGTGGTGCCGAGGTTCGGCTGGTGGAGTCTGGCGGTGGCCTGGTGCA<br>GCCAGGGGGCTCACTCCGTTTGTCCTGTGCAGCTTCTGGCTTCAACATCTACT<br>ATGGTTCTTACATCCACTGGGTGCGTCAGGCCCCGGGTAAGGGCCTGGAATG<br>GGTTGCAGGTATTTCTCCTTACTACAACTCTACTTACTATGCCGATAGCGTCAA<br>GGGCCGTTTCACTATAAGCGCAGACGCATCCAAAAACACAGCCTACCTACAAA<br>TGAACAGCTTAAGAGCTGAGGACACTGCCGTCTATTATTGTGCTCGCGGTTAC<br>GCTTCTGGTGGTATGGACTACTGGGGTCAAGGAACCCTGGTCACCGTCTCCT<br>CGGGATCC |

**Table S2.** Primary and secondary antibody labeling for flow cytometry characterization and FACS experiments.

| Detection | Primary Label (dilution) | Secondary Label (dilution) |
|---|---|---|
| **c-Myc epitope tag** | Chicken anti-cMyc (1:500) | Goat anti-chicken Alexa Fluor 647 (1:500) |
| **MMP-9 binding** | Mouse anti-His (1:500) | Goat anti-mouse Alexa Fluor 488 (1:500) |

**Table S3.** Number of cells, displayed constructs, effective scFv concentration (assuming $5\times10^4$ displayed constructs per cell) and concentration ratios used for titrating yeast displayed constructs for inhibition assays in 96-well plates.

| | Number of Cells | Number of displayed constructs | Effective concentration (nM) | Concentration ratio |
|---|---|---|---|---|
| **A1** | 0 | 0 | 0 | 0:1 |
| **B1** | 240,800 | $1.204\times10^{10}$ | 0.2 | 0.1:1 |
| **C1** | 2,408,000 | $1.204\times10^{11}$ | 2 | 1:1 |
| **D1** | 12,040,000 | $6.020\times10^{11}$ | 10 | 5:1 |
| **E1** | 24,080,000 | $1.204\times10^{12}$ | 20 | 10:1 |
| **F1** | 48,160,000 | $2.408\times10^{12}$ | 40 | 20:1 |
| **G1** | 120,400,000 | $6.020\times10^{12}$ | 100 | 50:1 |
| **H1** | 240,800,000 | $1.204\times10^{13}$ | 200 | 100:1 |

### Supplementary Figures

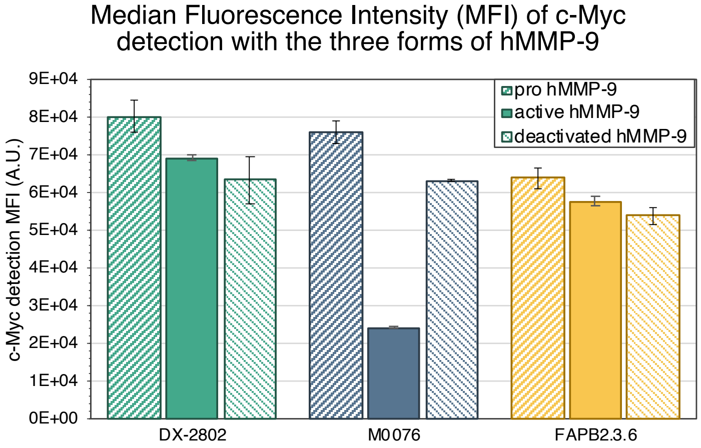


**Figure S1**. Median Fluorescence Intensity (MFI) of c-Myc detection for each of the three constructs when incubated with the three forms of hMMP-9 (pro hMMP-9, active hMMP-9 and EDTA treated hMMP-9)

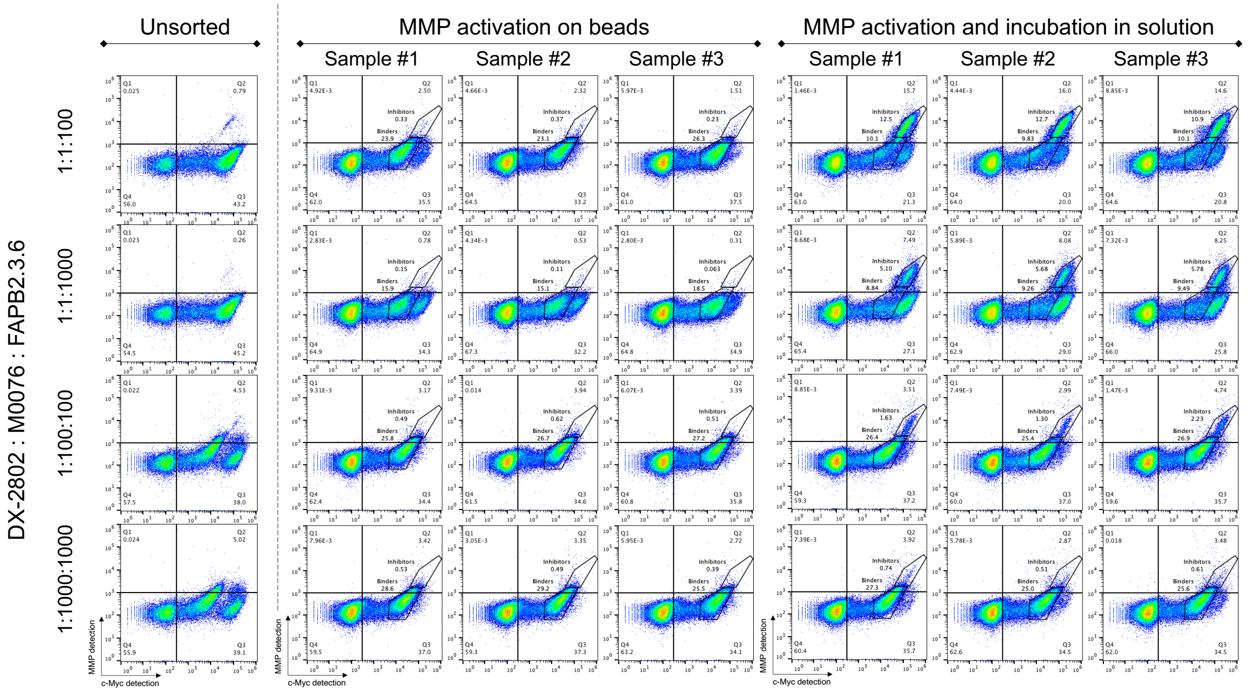


**Figure S2**. Magnetic bead-based enrichment results for four distinct mixture compositions of DX-2802, M0076 and FAPB2.3.6. Flow cytometry dot plots of unsorted populations (left), sorted with MMP activated on beads (middle) in triplicates, and sorted with MMP activated and incubated in solution (right) in triplicates. Gates indicate populations corresponding to DX-2802 (Inhibitors) and M0076 (Binders) with their corresponding abundance.

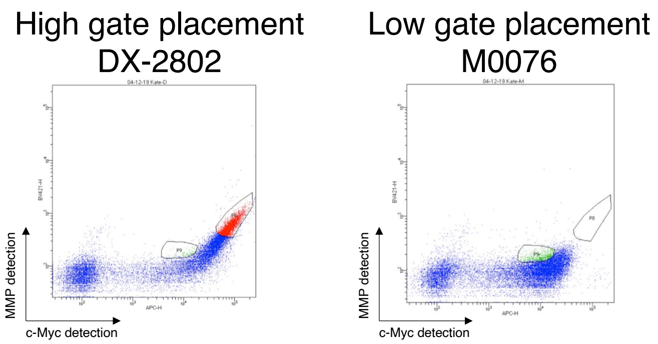


**Figure S3**. Gate placement prior to FACS sorting. High gate placement established from a single DX-2802 clone (left), low gate placement established from a single M0076 clone (right).

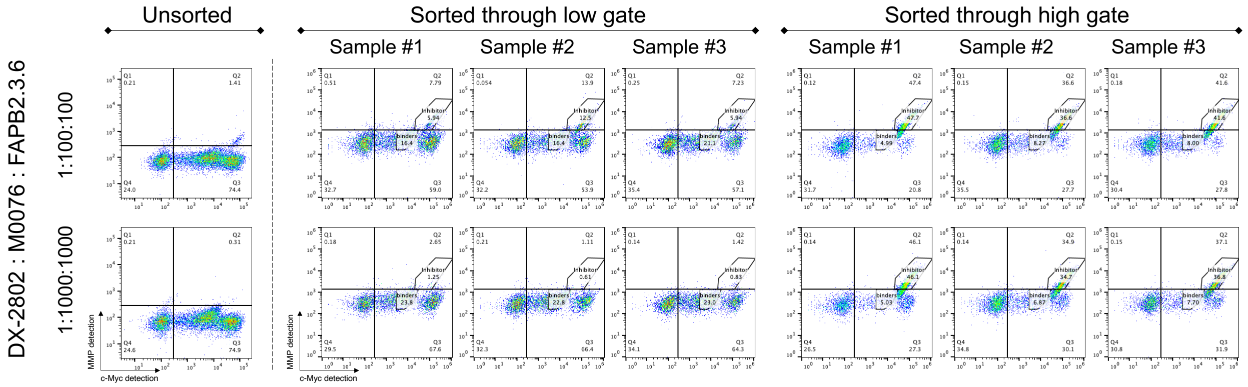


**Figure S4**. FACS enrichment results for two distinct mixture compositions of DX-2802, M0076 and FAPB2.3.6. Flow cytometry dot plots of unsorted populations (left), FACS sorted populations with a high gate (middle) in triplicates, and sorted with a low gate (right) in triplicates. Gates indicate populations corresponding to DX-2802 (Inhibitors) and M0076 (Binders) with their corresponding abundance.

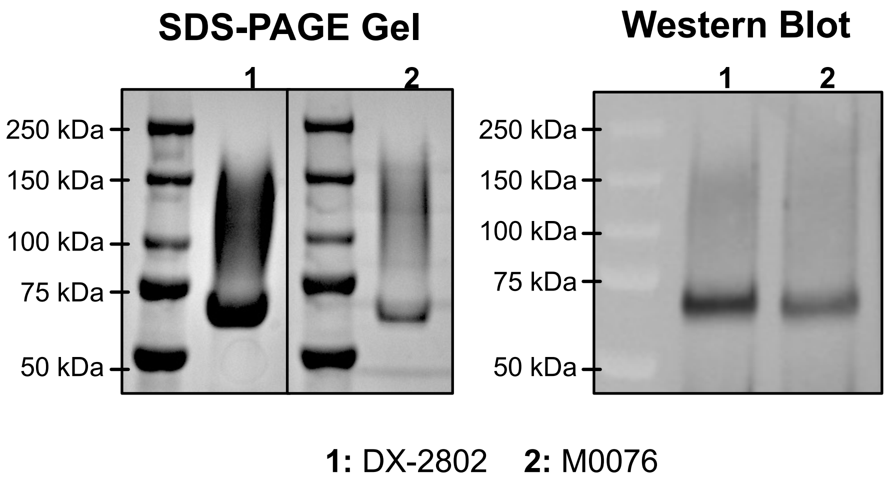


**Figure S5**. SDS-PAGE and Western Blot visualized gels following production of DX-2802 and M0076 constructs in soluble form.

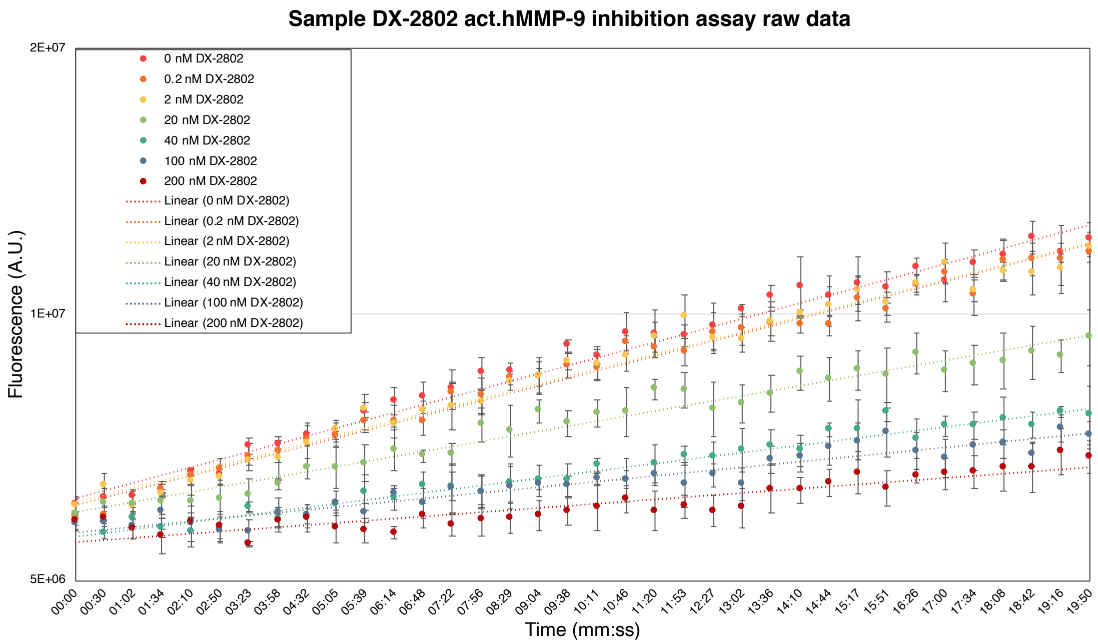


| Inhibitor Concentration (nM) | Calculated slope | | | Fractional activity | | |
|---|---|---|---|---|---|---|
| | Sample 1 | Sample 2 | Sample 3 | Sample 1 | Sample 2 | Sample 3 |
| **0** | 3.93E+08 | 3.29E+08 | 3.96E+08 | | | |
| **0.2** | 3.79E+08 | 3.33E+08 | 3.56E+08 | 0.96560 | 1.01465 | 0.89775 |
| **2** | 3.64E+08 | 3.26E+08 | 3.72E+08 | 0.92699 | 0.99352 | 0.93823 |
| **10** | 2.59E+08 | 2.05E+08 | 2.58E+08 | 0.66056 | 0.62346 | 0.65076 |
| **20** | 2.03E+08 | 1.71E+08 | 1.48E+08 | 0.51603 | 0.52038 | 0.37262 |
| **40** | 1.57E+08 | 1.06E+08 | 1.4E+08 | 0.40062 | 0.32331 | 0.35449 |
| **100** | 1.17E+08 | 69866159 | 1.15E+08 | 0.29725 | 0.21267 | 0.29126 |
| **200** | 85154143 | 78303878 | 79680038 | 0.21678 | 0.23836 | 0.20105 |
| | **Calculated slope at concentration $i = v_i$** | | | **Fractional activity at concentration $i = \frac{v_i}{v_0}$** | | |

THREE BIOLOGICAL REPLICATES

Normalized through excel formula below, then interpolated via GraphPad Prism v9.1 using Sigmoidal, 4PL parameters to obtain $IC_{50}$ values

Normalized fractional activity at concentration $i$ =

$$\frac{\text{Fractional Activity}_i - \text{MIN(Fractional acvity of sample)}}{\text{MAX(Fractional activity of sample)} - \text{MIN(Fractional activity of sample)}}$$

Normalized through GraphPad Prism v9.1, then interpolated using Sigmoidal, 4PL parameters to obtain $IC_{50}$ values.

MATCHING RESULTS

**Figure S6**. Step-by-step analysis of inhibition assay data to obtain fractional activity for each condition, normalize data and interpolate to obtain $IC_{50}$ values. Each technical triplicate was conducted in three biological replicates and the normalization process was conducted in two different ways (Excel formula on the left, normalization through prism on the right) to verify conclusions.
